# An unsupervised map of excitatory neurons’ dendritic morphology in the mouse visual cortex

**DOI:** 10.1101/2022.12.22.521541

**Authors:** Marissa A. Weis, Stelios Papadopoulos, Laura Hansel, Timo Lüddecke, Brendan Celii, Paul G. Fahey, Eric Y. Wang, J. Alexander Bae, Agnes L. Bodor, Derrick Brittain, JoAnn Buchanan, Daniel J. Bumbarger, Manuel A. Castro, Forrest Collman, Nuno Maçarico da Costa, Sven Dorkenwald, Leila Elabbady, Akhilesh Halageri, Zhen Jia, Chris Jordan, Dan Kapner, Nico Kemnitz, Sam Kinn, Kisuk Lee, Kai Li, Ran Lu, Thomas Macrina, Gayathri Mahalingam, Eric Mitchell, Shanka Subhra Mondal, Shang Mu, Barak Nehoran, Sergiy Popovych, R. Clay Reid, Casey M. Schneider-Mizell, H. Sebastian Seung, William Silversmith, Marc Takeno, Russel Torres, Nicholas L. Turner, William Wong, Jingpeng Wu, Wenjing Yin, Szi-chieh Yu, Jacob Reimer, Philipp Berens, Andreas S. Tolias, Alexander S. Ecker

## Abstract

Neurons in the neocortex exhibit astonishing morphological diversity which is critical for properly wiring neural circuits and giving neurons their functional properties. However, the organizational principles underlying this morphological diversity remain an open question. Here, we took a data-driven approach using graph-based machine learning methods to obtain a low-dimensional morphological “bar code” describing more than 30,000 excitatory neurons in mouse visual areas V1, AL and RL that were reconstructed from the millimeter scale MICrONS serial-section electron microscopy volume. Contrary to previous classifications into discrete morphological types (m-types), our data-driven approach suggests that the morphological landscape of cortical excitatory neurons is better described as a continuum, with a few notable exceptions in layers 5 and 6. Dendritic morphologies in layers 2–3 exhibited a trend towards a decreasing width of the dendritic arbor and a smaller tuft with increasing cortical depth. Inter-area differences were most evident in layer 4, where V1 contained more atufted neurons than higher visual areas. Moreover, we discovered neurons in V1 on the border to layer 5 which avoided deeper layers with their dendrites. In summary, we suggest that excitatory neurons’ morphological diversity is better understood by considering axes of variation than using distinct m-types.

## 1 Introduction

Neurons have incredibly complex and diverse shapes. Since Ramón y Cajal, neuroanatomists have studied their morphology [29] and have classified them into different types. From a computational point of view, a neuron’s dendritic morphology constrains which inputs it receives, how these inputs are integrated and, thus, which computations the neuron and the circuit it is part of can learn to perform.

Less than 15% of neocortical neurons are inhibitory, yet they are morphologically the most diverse and can be classified reliably into well-defined subtypes [7, 13]. The vast majority of cortical neurons are excitatory. Excitatory cells can be divided into spiny stellate and pyramidal cells [26]. Although pyramidal cells have a very stereotypical dendritic morphology, they exhibit a large degree of morphological diversity. Recent studies subdivide them into 10–20 cell types using manual classification [21] or clustering algorithms applied to dendritic morphological features [10, 16, 25].

Existing studies of excitatory morphologies have revealed a number of consistent patterns, such as the well-known thick-tufted pyramidal cells of layer 5 [14, 10, 16, 21, 25]. However, a commonly agreed-upon morphological taxonomy of excitatory neuron types is yet to be established. For instance, Markram et al. [21] describe two types of thick-tufted pyramidal cells based on the location of the bifurcation point of the apical dendrite (early vs. late). Later studies suggest that these form two ends of a continuous spectrum [16, 10]. Other authors even observe that morphological features overall do not form isolated clusters and suggest an organization into families with more continuous variation within families [31]. There are two main limitations of previous morphological characterizations: First, many rely on relatively small numbers of reconstructed neurons used to asses the morphological landscape. Second, they represent the dendritic morphology using summary statistics such as point counts, segment lengths, volumes, density profiles (so-called morphometrics; [25, 33, 20]) or graph-based topological measures [15]. These features were handcrafted by humans and may not capture all crucial axes of variation.

We here take a data-driven approach using a recently developed unsupervised representation learning approach [42] to extract a morphological feature representation directly from the dendritic skeleton. We apply this approach to a large-scale anatomical dataset [6] to obtain low-dimensional vector embeddings (“bar codes”) of more than 30,000 neurons in mouse visual areas V1, AL and RL. Our analysis suggests that excitatory neurons’ morphologies form a continuum, with notable exceptions such as layer 5 thick-tufted cells, and vary with respect to three major axes: soma depth, total apical and total basal skeletal length. Moreover, we found a number of novel morphological features in the upper layers: Neurons in layers 2/3 showed a trend of a decreasing width of their dendritic arbor and a smaller tuft with increasing cortical depth. In layer 4, morphologies showed area-specific variation: atufted neurons were primarily located in the primary visual cortex, while tufted neurons were more abundant in higher visual areas. Finally, layer 4 neurons in V1 on the border to layer 5 showed a tendency towards avoiding layer 5 with their dendrites.

## 2 Results

### 2.1 Self-supervised learning of embeddings for 30,000 excitatory neurons from visual cortex

Our goal was to perform a large-scale census of the dendritic morphologies of excitatory neurons without prescribing a-priori which morphological features to use. Therefore, we used machine learning techniques [42] to learn the features directly from the neuronal morphology.

Our starting point was a 1.3×0.87×0.82 mm^3^ volume of tissue from the visual cortex of an adult P75–87 mouse, which has been densely reconstructed using serial section electron microscopy [6]. This volume has been segmented into individual cells, including non-neuronal types and more than 54,000 neurons whose soma was located within the volume. From these detailed reconstructions we extracted each neuron’s dendritic tree and represented it as a skeleton (Fig. 1A) [4]: each neuron’s dendritic morphology was represented as a graph, where each node had a location in 3d space. This means we focused on the location and branching patterns of the dendritic tree, not fine-grained details of spines or synapses (see companion paper [32]), or any subcellular structures (see companion paper [8]).

**Figure 1:**
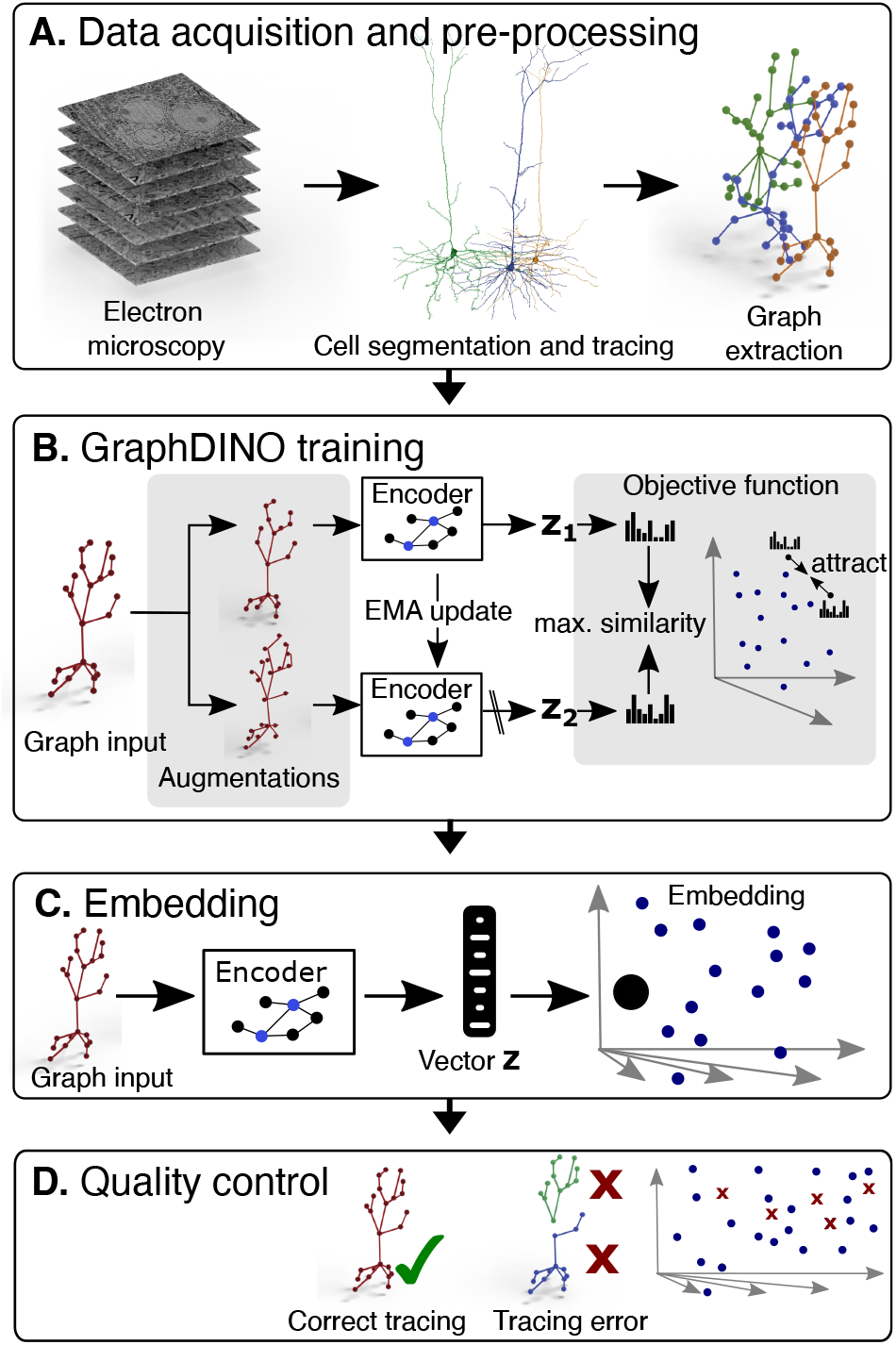
Pipeline to generate vector embeddings for large scale datasets that capture the morphological features of the neurons’ dendritic trees. **A**. Imaging of brain volume via electron microscopy and subsequent segmentation and tracing to render 3D meshes of individual neurons that are used for skeletonization. **B**. Self-supervised learning of low dimensional vector embeddings *z*1, *z*2 that capture the essence of the 3D morphology of individual neurons using Graphdino. Two augmented “views” of the neuron are input into the network, where the weights of one encoder (bottom) are an exponential moving average (EMA) of the other encoder (top). The objective is to maximize the similarity between the vector embeddings of both views. Vector embeddings of similar neurons are close to each other in latent space. **C**. An individual neuron is represented by its vector embedding as a point in the 32-dimensional vector space. **D**. Quality control to remove neurons with tracing errors.

Our next step was to embed these graphs into a vector space that defined a measure of similarity, such that similar morphologies were mapped onto nearby points in embedding space (Fig. 1B). To do so, we employed a recently developed self-supervised learning method called GraphDINO [42] that learns semantic representations of graphs without relying on manual annotations. The idea of this method is to generate two “views” of the same input by applying random identity-preserving transformations such as rotations around the vertical axis, slightly perturbing node locations or dropping subbranches (Fig. 1B, top and bottom). Then both views are encoded using a neural network. The neural network is trained to map both views onto similar vector embeddings. For model training, the data was split into training, validation and test data to ensure that the model did not overfit (Sec. 4.5). The model outputs a 32-dimensional vector for each neuron that captures the morphological features of the neuron’s dendritic tree. Thus, each neuron is represented as a point in this 32-dimensional vector space (Fig. 1C).

At this stage, we performed another quality control step: Using the learned embeddings as a similarity metric between neurons, we clustered the neurons into 100 clusters and manually inspected the resulting clusters. We found a non-negligible fraction of neurons whose apical dendrite left the volume or was lost during tracing (see Methods for details). We removed neurons whose somata are in close proximity to the imaged volume boundary (Fig. 2A). Additionally, we used the clusters containing fragmented neurons as examples for broken neurons and trained a classifier to predict whether a neuron has reconstruction errors using the learned morphological embeddings as input features (Fig. 2B, Fig. A.2A, B). We then removed all neurons from the dataset that were classified as erroneous. Also, at this point we removed all interneurons from the dataset since we focused on excitatory neurons in this paper (Fig. 2C, Fig. A.2B, C).

**Figure 2:**
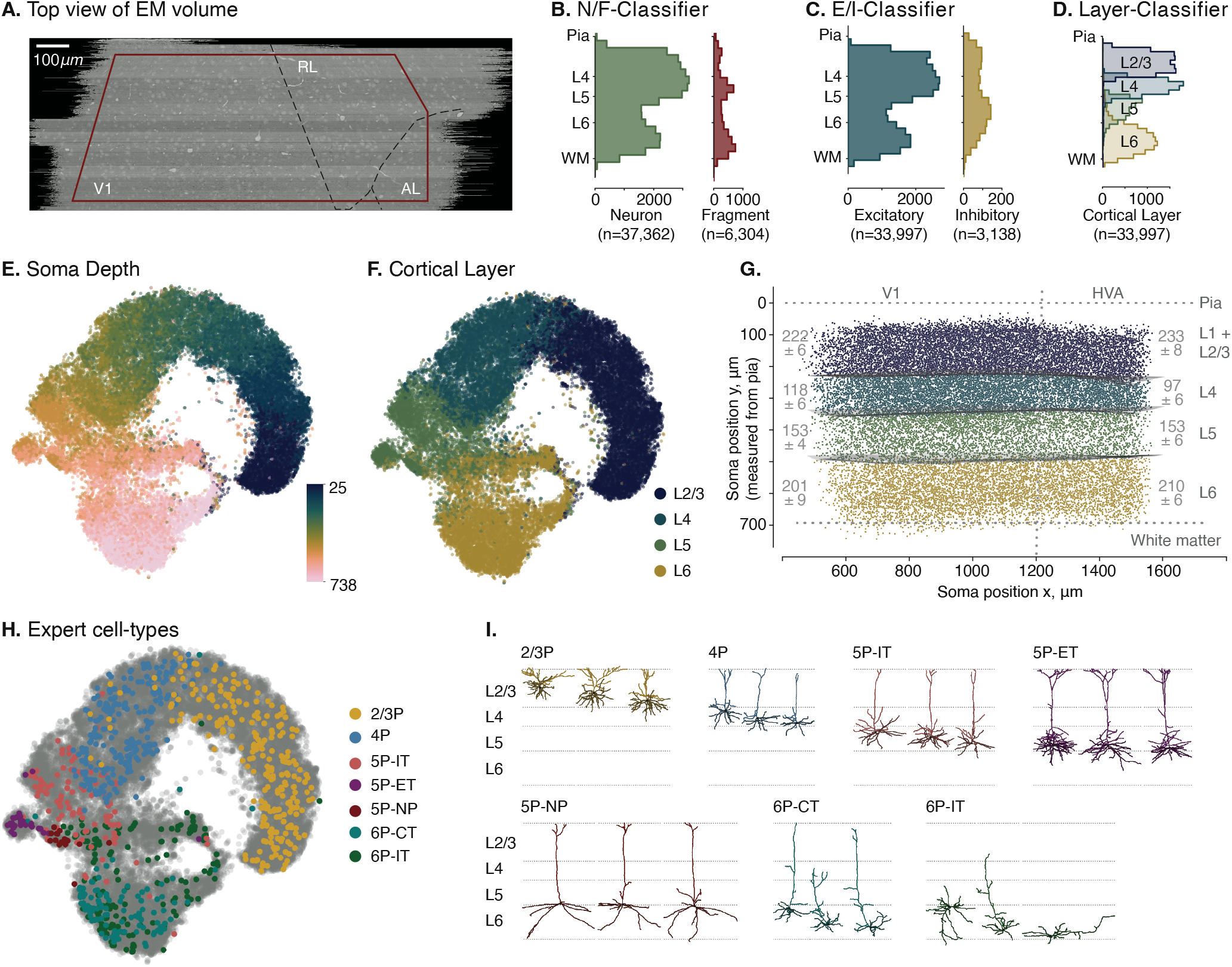
Visualization of soma depths and cortical layer assignments of excitatory neuronal morphologies showing mostly a continuum with distinct clusters only in deeper layers. **A**. Top view of the EM volume with approximate visual areas indicated. All neurons with their soma origin within the red boundary were used for analysis. **B**. Distribution of complete neurons and fragments along cortical depth as determined by our classifier based on the morphological embeddings. **C**. Distribution of excitatory neurons and interneurons along cortical depth. **D**. Classifier prediction for cortical layer origin based on the learned morphological embeddings. **E**. t-SNE embedding (perplexity = 300) of the vector embeddings of excitatory neuronal morphologies colored by the respective soma depth of the neurons relative to the pia (*n* = 33,997). **F**. t-SNE embedding colored by cortical layer assignments as predicted by a cross-validated classifier trained on the morphological embeddings as features and a subset of manually labeled excitatory neurons (*n* = 922). **G**. Cross-section of the brain volume depicting soma positions of neurons colored by their assigned cortical layer. Cortical layer thicknesses for primary visual cortex (V1) (left) and higher visual areas (HVA) (right) given as mean ± standard deviation. **H**. t-SNE embedding of excitatory neuronal morphologies colored by expert-defined cell types. **I**. Example morphologies of the expert-defined cell types.

The vector embeddings of the remaining 33,997 excitatory neurons in the dataset were organized by cortical depth (Fig. 2E) and, as a consequence, could distinguish well between different cortical layers (Fig. 2F, G; note that there is no 1:1 correspondence between cortical depth and layer as the layer boundaries varied across the volume.). The learned embeddings could also distinguish between broad cell types (Fig. 2H, I) that were assigned by expert neuroanatomists [32] using cortical origin of the somata and their long-range projection type (IT: intratelencephalic or intracortical; ET: extratelencephalic or subcortical projecting, NP: near projecting, and CT: cortico-thalamic). Note that neither the location of the soma nor the projection type were provided to the model, showing that the dendritic morphology by itself provides information on these broad cell types. One exception are the 6P-CT and 6P-IT cells, which were partly intermingled in embedding space. 6P-IT cells show a high variance in their dendritic morphology which in some cases are indistinguishable from 6P-CT cells when no information about the projection type is used (Fig. 2H, I).

To test that the learned embedding is generally applicable beyond EM datasets and the MICrONS dataset specifically, we used the GraphDINO model trained on MICrONS to embed 61 neurons from mouse visual cortex [1] that have been recorded using PatchSeq [2] and show that the model generalizes to other datasets and recording techniques (Fig. A.10).

### 2.2 Dendritic morphologies mostly form a continuum with distinct clusters only in deeper layers

We noticed that the embedding space appeared to form largely a continuum, with only a few fairly distinct clusters, such as the layer 5 ET cells (Fig. 2H, purple). Previous papers have characterized excitatory morphologies by categorizing neurons into morphological types (m-types), with the number of types varying between nine and nineteen [25, 21, 24, 15, 10, 32]. But is categorization into discrete types the best way of describing the landscape of morphologies or is it rather characterized by continuous variation? The answer depends on the structure of the data. Consider the following toy example where the data is generated by a mixture of two normal distributions (Fig. 3A): If the two components are well separated, it makes sense to define each one as a distinct type (Fig. 3A, left). However, if they are strongly overlapping such that the resulting data distribution is not even bimodal (Fig. 3A, right), describing the distribution by two types is not useful and identifying the two types by clustering will not work reliably, either. But there are also scenarios in between, where the distinction is not as straightforward (Fig. 3A, middle). Thus, the question of whether a distribution is discrete or forms a continuum does not have a binary answer – it is rather a matter of degree.

**Figure 3:**
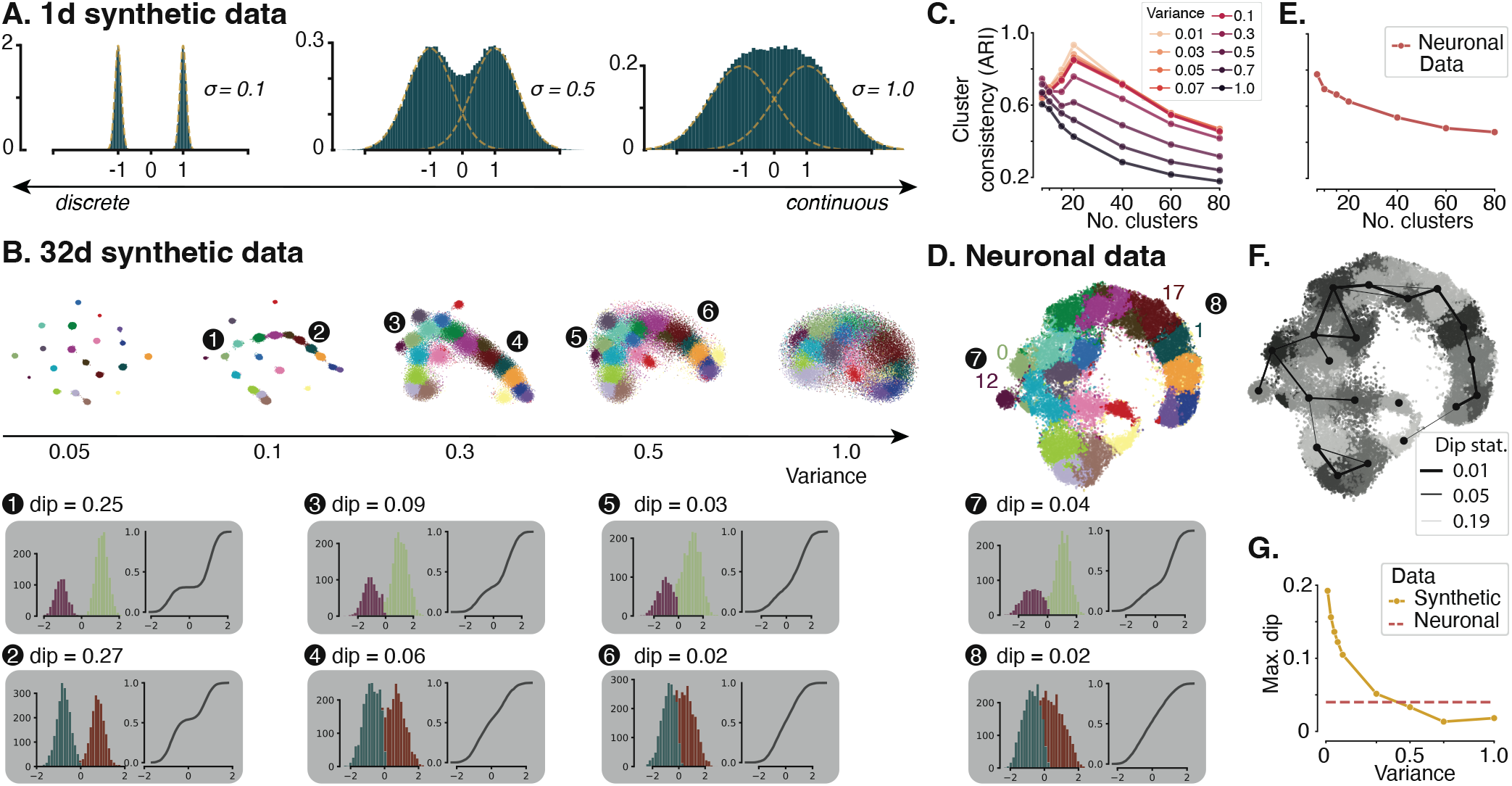
Cluster versus continuum analysis: **A**. Histograms of samples from a 1d Gaussian mixture (*n* = 30 000, number of components = 2) in green and the underlying mixture components with means *µ*1 = −1 and *µ*2 = 1 in yellow. Data distributions evolve from discrete to continuous by increasing the standard deviation (SD) from left to right. **B**. t-SNE representation of synthetic data (*n* = 33 997, perplexity= 300). Synthetic data is sampled from Gaussian mixtures with 20 components. Cluster means and weights are estimated from neuronal data. Isotropic variance is set to obtain data evolving from discrete clusters to uniform distributions. Grey insets (1–6) show histograms of two sample clusters (12 and 1) and their nearest neighbors (0 and 17, respectively) projected onto the direction connecting their cluster means (left), as well as the cumulative distribution of the samples assigned to these two clusters’ along this direction (right). **C**. Mean adjusted rand index (ARI) of 100 GMMs with increasing number of components fit to the synthetic datasets. The correct number of underlying components can be identified as long as the variance in the data is not too high (< 0.5 for 20 components). **D**. t-SNE representation (*n* = 33 997, perplexity= 300) of neuronal data colored by cluster membership (GMM with 20 components). Grey insets (7 & 8) show 1d projections of the clusters 12 and 1 onto the line connecting their means with their nearest neighbors (0 and 17, respectively). Cumulative distributions show that while there is a gap between cluster 12 and its neighbors, there is none between cluster 1 and its neighbors. **E**. Cluster analysis as in **C**. for neuronal data. No specific number of components can be recovered. **F**. t-SNE representation of neuronal data overlaid with nearest neighbor graph between clusters. Line width indicates dip statistic (thicker = more connected). **G**. Maximum dip statistic between all clusters and their nearest neighbor for the synthetic data with 20 components and varying variance (yellow curve) and for the neuronal data clustered with 20 components (red dashed line).

To understand to what degree our dataset forms a continuum, we devised a simple procedure based on synthetic data that emulates the real data to some extent but allows us to manipulate the degree of separation. The synthetic data was generated from a Gaussian mixture model (GMM) fit to our morphological embeddings, from which we kept the cluster means and weights, but replaced the covariance matrix to be spherical with varying variances (*σ*^2^). Following previous estimations of number of excitatory cell types in the rodent sensory cortex [25, 21, 24, 15, 10, 32], we generated synthetic data distributions with 20 clusters (Fig. 3B), as well as with 10 and 40 clusters as controls (Fig. A.5). When the variance was small (*σ*^2^ = 0.05), all clusters were clearly distinct (Fig. 3B, left). As we increased the variance to 1, the distribution became more and more continuous (Fig. 3B, right). At intermediate values of 0.3 or 0.5 the synthetic data distribution resembled qualitatively the real data (Fig. 3D).

To make the comparison more quantitative, we asked two questions, which can be answered using the synthetic data for which the ground truth generating process is known. First, we asked under which conditions we could reliably identify the underlying clusters that generated the data (Fig. 3C). To do so, we assumed we did not know the generative process and clustered the synthetic data repeatedly by fitting Gaussian mixture models (GMMs) with varying number of components and random initial conditions. We found that in the extreme scenario when all clusters were clearly separated, the result of the clustering was highly consistent across runs when the number of clusters matched the ground truth (Fig. 3C; ARI ≥ 0.85 for *σ*^2^ ≤ 0.1 and number of ground truth components equal to number of GMM components). As the degree of overlap between the clusters increased, the consistency of the clustering result decreased and the optimal number of clusters was increasingly less clearly defined. For a larger degree of overlap (*σ*^2^ *>* 0.5) the consistency of clusterings decreased monotonically with the number of clusters and no optimal number of clusters could be determined. The same was true for the real data (Fig. 3E): There was no noticeable peak in the ARI across different number of clusters, suggesting that the scenario with *σ*^2^ ≥ 0.5 is realistic in this regard (Neuronal data: ARI = 0.63 for number of clusters = 20; compared to ARI = 0.62 for number of clusters = 20 and *σ*^2^ = 0.5 for the synthetic data).

Next, we investigated the degree to which individual clusters were distinct from their neighboring clusters. Even though certain parts of the distribution appeared continuous, there could be clusters that are separable. To address this question, we built a *k*-nearest-neighbor graph from the clustering output, connecting each cluster to its *k* =3 nearest neighbors. We then quantified for each pair of neighboring clusters how separated they are. To do so, we projected all data points assigned to the pair onto the direction connecting the two cluster means (Fig. 3B, insets left) and computed the dip statistic [12]. The dip statistic measures how bimodal a distribution is by computing how much its empirical cumulative distribution deviates from that of the closest uniform distribution (Fig. 3B, insets right). It is close to zero for unimodal distributions and increases with increasing separation of the two modes of a bimodal distribution. This analysis confirmed the qualitative impression from the t-distributed stochastic neighbor embedding (t-SNE; [37]) that the layer 5 ET cluster (purple cluster 12 in Fig. 3B, D) was separated more from its nearest neighbor (cluster 0, green) than two representative example clusters from layer 2/3 (clusters 1 and 17, red and teal), which were not separated and appeared to divide a continuum more or less arbitrarily. These two patterns of results in the neuronal data were reproduced well by the synthetic data with a standard deviation of 0.5 (Fig. 3B, insets 5 & 6). Examination of the entire nearest-neighbor graph showed that layers 2–4, including the upper part of layer 5, form a continuum with no neighboring clusters being well-separated, clusters in layer 5 were more distinct, and two clusters in layer 6 (inverted and subplate neurons) stood out from a larger clique of layer 6 clusters (Fig. 3F). Over the entire dataset, the maximum dip statistic (maximally separated clusters) of the neuronal data was in between the maximal dip statistic for the synthetic data with *σ*^2^ = 0.3 and *σ*^2^ = 0.5 (Fig. 3G), again suggesting that the qualitative visualization by t-SNE captures the underlying structure of the data well.

The analyses presented so far established that our learned morphological embeddings form mostly a continuum. Could this result be caused by our learning methods? This was not the case, as using a different contrastive learning objective (Fig. A.7) to train GraphDINO and varying model hyperparameters (Fig. A.6) produced the same result. Similarly, using handcrafted morphometrics from earlier studies [10, 32] on our data did not change our conclusions (Fig. A.9, Sec. A.1). Additionally, we employed alternative dimensionality reduction techniques with varying settings (Fig. A.8) to ensure that our interpretation is not dependent on t-SNE for visualization.

### 2.3 The landscape of morphological variation across layers

Given the results from the previous section, we conclude that excitatory morphologies were mostly organized along a continuum, with only a few distinct clusters in the deeper layers. Therefore we did not base our subsequent analyses on a set of m-types as previous studies did, but instead investigated the major axes of variation within the morphological embedding space. The cortical organization into layers is well established, so we separated cells by cortical layer. We determined the layer boundaries by training a classifier using our 32-dimensional morphological embeddings and a set of 922 neurons manually assigned to layers by experts (Fig. 2D, F, G). As expected, the inferred layer boundaries indicated that layer 4 was approximately 20% thicker in V1 than in higher visual areas RL and AL (Fig. 2G; mean ± SD: 118 ± 6 µm in V1 vs. 97 ± 6 µm in HVA), the difference being compensated for by layers 2/3 and 6 each being approximately 10 µm thinner. In the following we proceed by assigning neurons to layers based on their soma location relative to these inferred boundaries.

To visualize the main axes of morphological variation within each layer, we performed nonlinear dimensionality reduction using t-SNE and identified morphological features that formed major axes of variation within the two-dimensional space (Fig. 5).

What do these axes of variation in the two-dimensional t-SNE embeddings mean in human-interpretable terms? To answer this question, we looked for morphological metrics that formed gradients within the t-SNE embedding space. Based on visual inspection, we found the following six morphological metrics to account well for a large fraction of the dendritic morphological diversity in our dataset (see Fig. 4 for an illustration): (1) depth of the soma relative to the pia, (2) height of the cell, (3) total length of the apical dendrites, (4) width of the apical dendritic tree, (5) total length of the basal dendrites, and (6) location of the basal dendritic tree relative to the soma (“basal bias”).

**Figure 4:**
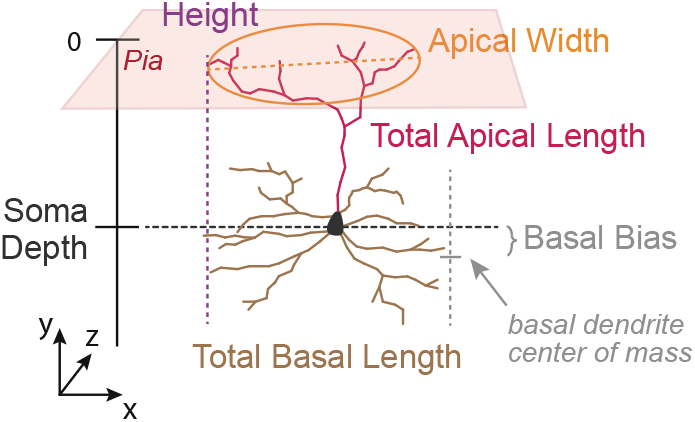
Schematic of morphometric descriptors computed from neuronal skeletons and their labeled compartments. SOMA DEPTH. Depth of the centroid of the soma relative to the pia. HEIGHT. Extent of the cell in y-axis. TOTAL APICAL LENGTH. Total length of the skeletal branches of the apical dendrites. APICAL WIDTH. Maximum extent of the apical dendritic tree in the xz-plane. TOTAL BASAL LENGTH. Total length of the skeletal branches of the basal dendrites. BASAL BIAS. Depth in y-axis of center of mass of basal dendrites relative to the soma.

### 2.4 Layer 2/3: Width and length of apical dendrites decrease with depth

In layer 2/3 (L2/3), we found a continuum of dendritic morphologies that formed a gradient from superficial to deep, with deeper neurons (in terms of soma depth) becoming thinner and less tufted (Fig. 5A L2/3 a,b,c). The strongest predictors of the embeddings were the depth of the soma relative to the pia and the total height of the cell (coefficient of determination *R*^2^ *>* 0.9; Fig. 5B L2/3; Tab. A.3). These two metrics were also strongly correlated (Spearman’s rank correlation coefficient, *ρ* = 0.93; Fig. 5C L2/3; Tab. A.4), since nearly all L2/3 cells had an apical dendritic tree that reached to the pial surface (see example morphologies in Fig. 5A L2/3, top). L2/3 cells varied in terms of their degree of tuftedness: both total length and width of their apical tuft decreased with the depth of the soma relative to the pia (Fig. 5A L2/3 b,c). L2/3 cells also varied along a third axis: the skeletal length of their basal dendrites (Fig. 5A L2/3 d), but this property was not strongly correlated with either soma depth or shape of the apical dendrites (Fig. 5C L2/3).

**Figure 5:**
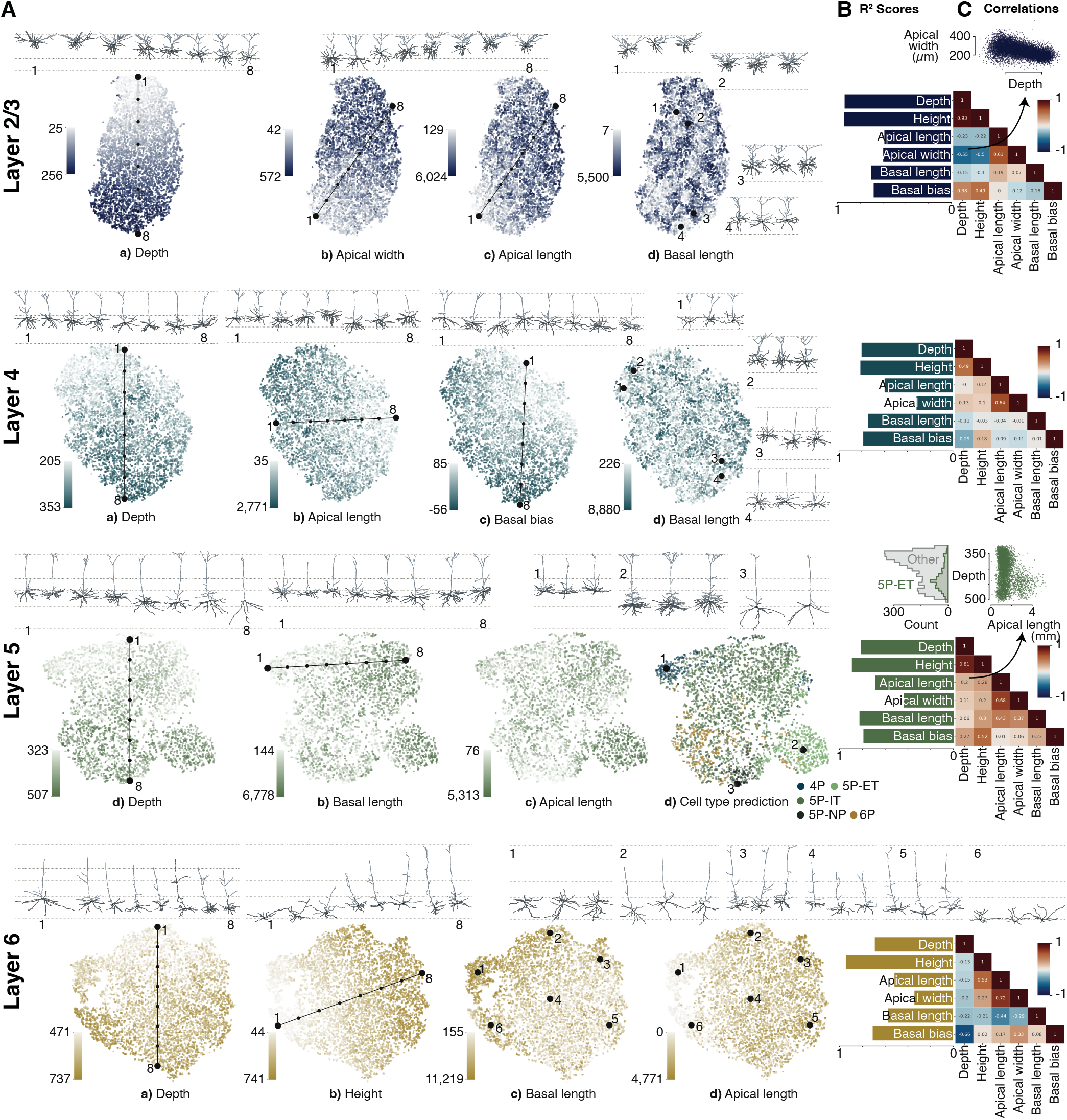
t-SNE visualization of vector embeddings per cortical layer reveal axes of variation in neuronal morphologies. **A**. t-SNE embeddings per layer colored by percentiles of various morphometric descriptors with example neuronal morphologies along the axis of variation displayed above the embedding. **B**. *R*^2^ scores of the six morphometric descriptors (see Fig. 4) per layer showing the strength as predictors of the 32d embeddings. **C**. Spearman’s rank correlation coefficient between morphometric descriptors per layer. **Layer 2/3** (blue) Continuum of dendritic morphologies with thinner and less tufted neurons in increasing distance to the pia. **Layer 4** (turquoise) Continuation of L2/3 trends with shorter apical dendrites and more atufted cells. Many cells avoid reaching dendrites into L5 (basal bias). **Layer 5** (green) Clustering of thick-tufted ET and NP cells. Upper L5 cells resemble L4 cells that avoid reaching into L5, indicating too strict laminar borders. **Layer 6** (orange) Continuum with a large morphological diversity e.g. in cell heights, and existence of horizontal and inverted pyramidal neurons.

### 2.5 Layer 4: Small or no tufts and some cells’ basal dendrites avoid layer 5

The dendritic morphology of layer 4 (L4) was again mostly a continuum and appeared to be a continuation of the trends from L2/3: The skeletal length of the apical dendrites was shorter, on average, than that of most L2/3 cells (Fig. 5A L4 b) and approximately 20% of the cells were atufted. Within L4 the total apical skeletal length was not correlated with the depth of the soma (*ρ* = 0.0; Fig. 5C L4; Tab. A.4), suggesting that it forms an independent axis of variation. Considerable variability was observed in terms of the total length of the basal dendritic tree, but – as in L2/3 – it was not correlated with any of the other properties.

Our data-driven embeddings revealed another axis of variation that had previously not been considered important: the location of the basal dendritic tree relative to the soma (“basal bias”; Fig. 4). We found that many L4 cells avoided reaching into L5 with their dendrites (Fig. 5A L4 c). As a result, the depth of the basal dendrites was anticorrelated with the depth of the soma (*ρ* = *-*0.29; Fig. 5A L4 c and Fig. 5C L4; Tab. A.4). We will come back to this observation later (see Sec. 2.9).

### 2.6 Layer 5: Thick-tufted cells stand out

Layer 5 (L5) showed a less uniformly distributed latent space than L2/3 or L4 (Fig. 5A L5, Fig. 3F). Most distinct was the cluster of well-known thick-tufted pyramidal tract (PT) cells [14, 10, 16, 21, 25] on the bottom right (Fig. 5A L5 d, light green points), also known as extratelencephalic (ET) projection neurons. These cells accounted for approximately 17% of the cells within L5 (based on a classifier trained on a smaller, manually annotated subset of the data; see Methods). They were restricted almost exclusively to the deeper half of L5 (Fig. 5A L5 a and d, inset 2; C inset top right) and compared to other L5 cells they have the longest skeleton for all three dendritic compartments: apical, basal and oblique.

Another morphologically distinct type of cell was apparent at the end of the layer 5 spectrum: the near-projecting (NP) cells [17, 10] with their long and sparse basal dendrites (Fig. 5A L5 d, inset 3). These cells accounted for approximately 4% of the cells within L5. They tended to send their dendrites deeper (relative to the soma), had little or no obliques and tended to have small or no apical tufts.

The remaining roughly 80% of the cells within L5 varied continuously in terms of the skeletal length of the different dendritic compartments. While there was a correlation between apical and basal skeletal length (apical vs. basal: *ρ* = 0.43; Fig. 5 L5 C; Tab. A.4), there was also a significant degree of diversity. Within this group there was no strong correlation of morphological features with the location of the soma within L5 (depth vs. apical length *ρ* = 0.2, depth vs. basal *ρ* = 0.06; Fig. 5 L5 C; Tab. A.4).

In upper L5 we found a group of cells that resembled the L4 cells whose dendrites avoid L5 (Fig. 5A L5 d, inset 1). These cells were restricted to the uppermost portion of L5 and morphologically resembled L4 cells by being mostly atufted and exhibiting upwards curved basal dendrites. We refer to these cells as displaced L4 cells. Their presence could be caused by our piece-wise linear estimation of the layer boundaries being not precise enough. Alternatively, it could suggest that there are no precise laminar boundaries based on morphological features of neurons, but instead different layers blend into one another as observed by previous studies [25, 8].

### 2.7 Layer 6: Long and narrow, oblique and inverted pyramidal neurons

Dendritic morphology in layer 6 (L6) also formed a continuum with a large degree of morphological diversity. The dominant feature of L6 was the large variety of cell heights (*R*^2^ *>* 0.9; Fig. 5 L6 B; Tab. A.3). Overall, the height of a cell was not strongly correlated with its soma’s location within L6 (*ρ* = −0.13; Fig. 5 L6 C; Tab. A.4). Unlike other layers, where the apical dendrites usually reach all the way up to layer 1, many cells in L6 have shorter apical dendrites. However, due to tracing errors, our analysis overestimated the number of such short cells. We therefore manually inspected 183 putative atufted early-terminating neurons within L6 and found that, among those, 45% were incompletely traced, whereas 55% were true atufted cells whose apical dendrite terminated clearly below L1 (Sec. 4.9).

As described previously [10, 25], the dendritic tree of L6 cells is narrower than in the layers above. Also consistent with previous work, we found a substantial number of horizontal and inverted pyramidal neurons, where the apical dendrite points sideways or downwards, respectively (Fig. 5A L6 d, inset 1 & 6). However, apicals of inverted and horizontal cells are currently not detected by the automatic compartment identification (see companion paper [4]), rendering an automatic analysis of the apical dendrites in layer 6 currently unreliable. This does not affect the learned embeddings as GraphDINO is trained without knowledge about the differentiation of dendritic compartments.

### 2.8 Pyramidal neurons are less tufted in V1 than in higher visual areas

After our layer-wise survey of excitatory neurons’ morphological features, we next asked whether there are inter-areal differences between primary visual cortex (V1) and higher visual areas (HVAs). The total length of the apical dendrites of neurons in V1 was significantly shorter than for neurons in HVA (Fig. 6A): For L2/3, neurons in V1 had on average 16% shorter apical branches than in HVA (mean ± SD: 1,423 ± 440 µm in V1 vs. 1,688 ± 554 µm in HVA; *t-test*: *p* < 10^−10^, Cohen’s *d* = 0.53). Similarly, L4 neurons in V1 had on average 16% shorter apical branches than in HVA (851 ± 264 µm vs. 1,019 ± 313 µm; *p* < 10^−10^, *d* = 0.58). In L5, neurons in V1 had on average 14% shorter apical branches than L5 neurons in HVA (1,326 ± 661 µm vs. 1,549 ± 745 µm; *p* < 10^−10^, *d* = 0.32). While the trend continued in L6, the difference in apical length between V1 and HVA neurons was smaller. There was only a 4% increase in apical length in HVA compared to V1 (1,112 ± 383 µm vs. 1,159 ± 397 µm; *p* = 1.810^−6^, *d* = 0.12). For this analysis, only neurons with identified apical dendrites were taken into account (see companion paper [4]).

**Figure 6:**
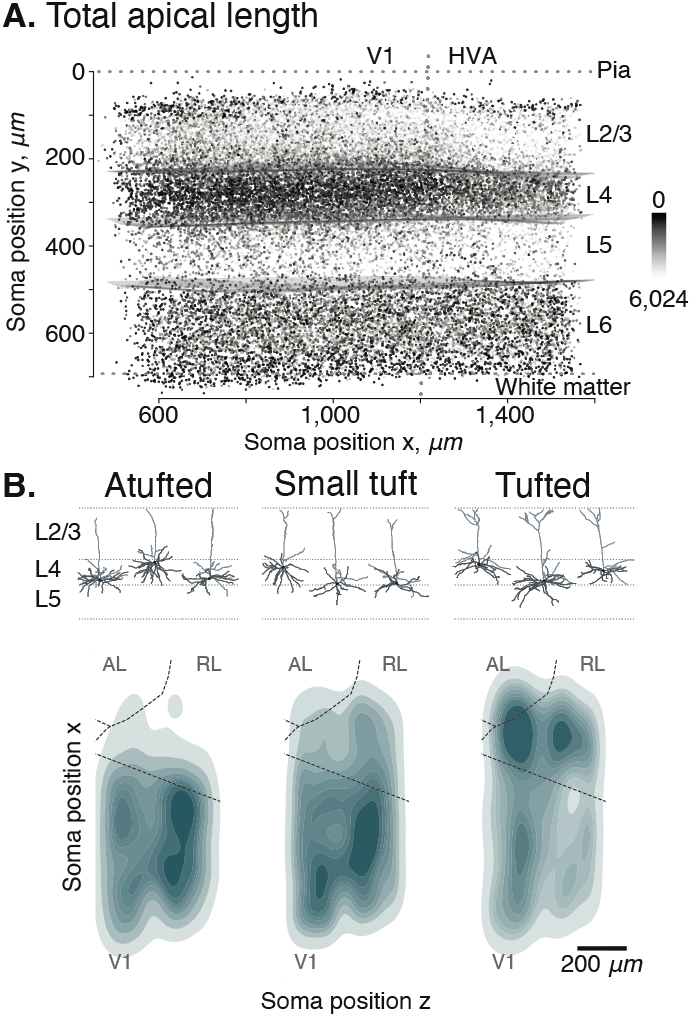
Inter-areal differences between primary visual cortex (V1) and higher visual areas (HVAs). **A**. Side view of the cortical volume. Each point represents the soma location of one neuron and is colored by apical skeletal length of the respective neuron (dark = no apical, bright = maximal apical skeleton length). Projection from the side orthogonal to the V1/HVA border after a 14 degree rotation around y-axis (vertical dashed line); top: pia; bottom: white matter. **B**. Top view of the volume showing the density of atufted (left), small tufted (middle) and tufted (right) L4 cells. Atufted neurons are mostly confined to V1, while tufted neurons are more abundant in HVA. Dashed lines: area borders between primary visual cortex (V1), anterolateral area (AL) and rostrolateral area (RL), estimated from reversal of the retinotopic map measured using functional imaging [6].

Upon closer inspection, we observed that L4 contained substantially more atufted neurons in V1 than in higher visual areas RL and AL (Fig. 6A). We clustered each layer’s morphological embeddings into 15 clusters using a Gaussian Mixture Model and looked for clusters that were restricted to particular brain areas. Clusters that were clearly confined to V1 or HVAs were primarily found in L4. When classifying (manually, at the cluster-level) L4 neurons into atufted, small tufted and tufted, we observed that atufted neurons were almost exclusively located in V1, while tufted neurons were more frequent in HVAs (Fig. 6B).

### 2.9 Layer 4 cells avoiding layer 5 are located primarily in primary visual cortex

We observed a second area difference which was related to the novel morphological cell type in L4. Recall that these cells’ dendrites avoid reaching into L5. Interestingly, these cells were located in a very narrow strip of approximately 50 µm above the border between L4 and L5 (Fig. 7A). Moreover, they were atufted and almost exclusively located in V1 (Fig. 7B).

**Figure 7:**
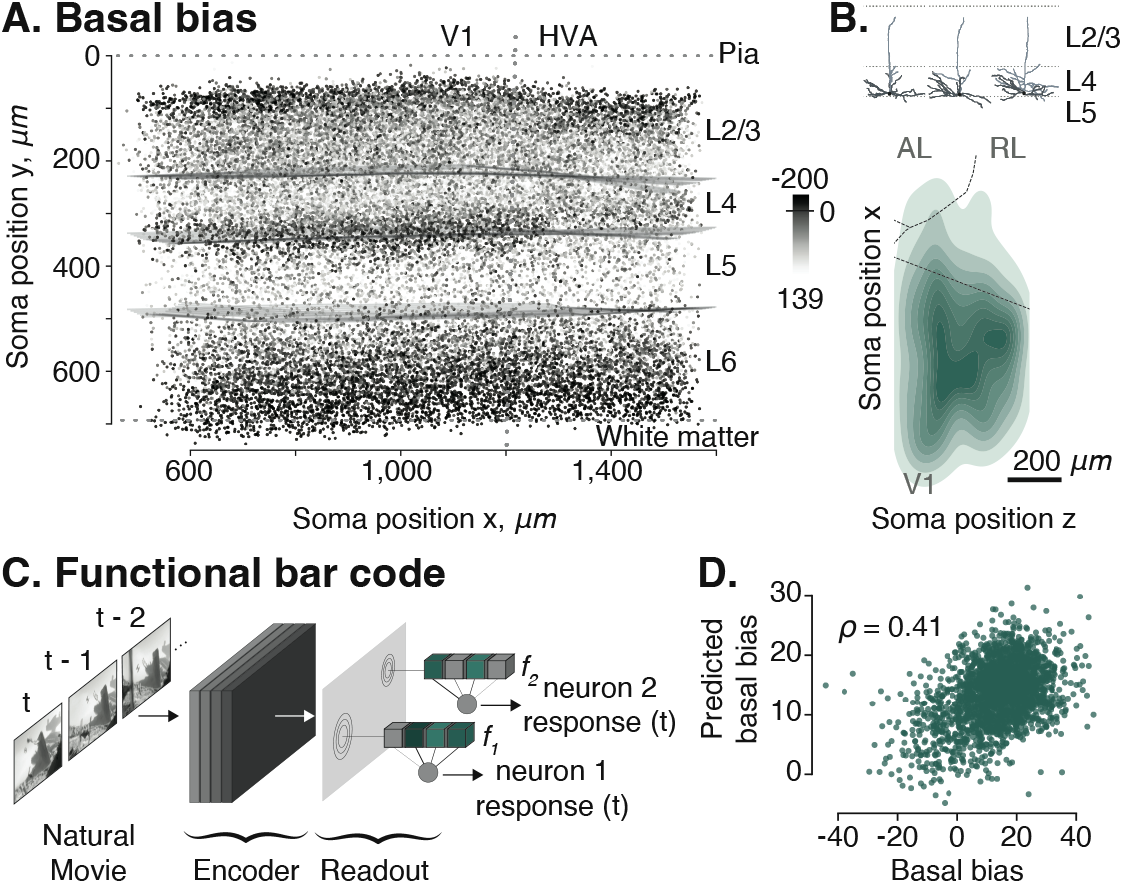
Basal bias neurons in primary visual cortex (V1). **A**. Side view of the cortical volume. Each point represents the soma location of one neuron and is colored by its respective basal bias (dark = negative basal bias: center of mass of basal dendrites is above the soma; bright = positive basal bias: center of mass of basal dendrites is below soma). **B**. Example neuronal morphologies of basal bias neurons (top) and top view of the volume (as in Fig. 6B) showing horizontal density distribution of L4 cells whose dendrites avoid reaching into L5 and who are mostly located in V1 (bottom). **C**. Functional digital twin can predict the functional response of the neurons to input stimuli such as natural movies. The input-output function of each neurons is described by a functional bar code *f*_*i*_ [38]. Schematic adapted from Consortium et al. [6] (Fig. 1). **D**. Predictions of basal bias metric from functional bar code *f*_*i*_ using linear regression.

### 2.10 Morphological property of avoiding layer 5 has functional correlate

Lastly, we asked whether morphological variation can be linked to the neurons’ functional properties. While an extensive investigation of the structure–function relationship is beyond the scope of this study, we took one novel morphological aspect revealed by our study as a proof of principle: We investigated whether L4 neurons that avoided reaching into layer 5 with their dendrites differ in their tuning to visual stimuli from other neurons in layer 4. To address this question, we made use of the fact that for many of the neurons in our dataset, we have measurements of how they respond to natural stimuli [6]. We leveraged a functional digital twin – a model that accurately predicted the response of a neuron to arbitrary visual stimuli [38] – to extract a functional bar code – a vector embedding *f*_*i*_ that describes the input-output function of a neuron analogous to how our morphological bar codes describe their morphology (Fig. 7C). From this functional bar code of each neuron, we predicted one of its morphological properties: the basal bias metric. We found that the basal bias of L4 neurons could be predicted reasonably well from the neurons’ response functions to visual stimuli (Fig. 7D; Pearson correlation *ρ* = 0.41, *p* < 10^−10^). This analysis could be confounded by cortical depth being predictive of the basal bias. However, a model predicting the basal bias from cortical depth and functional bar code explained significantly more variance in the basal bias metric than one using only cortical depth as predictor (*R*^2^ = 0.28 for both predictors vs. 0.21 for depth only; *ρ* = 0.53 and *ρ* = 0.46, respectively; Fisher’s z-test of difference between the correlation coefficients: *p* = 0.0015).

## 3. Discussion

In summary, our data-driven unsupervised learning approach identified the known morphological features of excitatory cortical neurons’ dendrites and enabled us to make four novel observations: (1) Superficial L2/3 neurons are wider than deep ones; (2) L4 neurons in V1 are less tufted than those in HVAs; (3) a novel atufted L4 cell type that is specific to V1 whose basal dendrites avoid reaching into L5; (4) excitatory cortical neurons form mostly a continuum with respect to dendritic morphology, with some notable exceptions.

First, our finding that superficial L2/3 neurons are wider than deeper ones is clearly visible in the data both qualitatively and quantitatively. A similar observation has been made recently in concurrent work [41].

Second, in L4 a substantial number of cells are completely atufted. Here we see a differentiation with respect to brain areas: completely atufted cells are mostly restricted to V1 while HVA neurons in L4 tend to be more tufted. Why would V1 neurons be less tufted than those in higher visual areas? V1 – as the first cortical area for visual information processing – and L4 – as the input layer, in particular – might be less modulated by feedback connections than other layers and higher visual areas. Therefore, these neurons might sample the feedback input in L1 less than other neurons.

Third, we found that some neurons at the bottom of L4 of V1 avoid reaching into L5 with their dendrites. To our knowledge, this morphological pattern has not been described before in the visual cortex. Retrospectively, it can be observed in Gouwens and colleagues’ data: their spiny m-types 4 and 5, which are smallor atufted L4 neurons, show a positive basal bias (assuming their “basal bias y” describes the same property; Gouwens et al. [10]; Suppl. Fig. 15). Whether such cells are restricted to the bottom of layer 4 or are simply morphologically insdistinguishable from other cells when located more superficially cannot be answered from our data. However, interestingly, this morphological pattern correlated with the functional properties of the neurons. While this is by no means an exhaustive characterization of *how* morphology and function are related, this result shows that they are, and that such relationships can be identified by data-driven methods. What function could avoiding L5 have? Similarly to the non-existing tuft of these neurons, avoiding L5 could support these neurons in focusing on the thalamic input (which targets primarily L4) and, thus, represent and distribute the feedforward drive within the local circuit. It is therefore tempting to speculate that these atufted, L5-avoiding L4 neurons might be precursors of spiny stellate cells, which are nearly absent in the mouse visual cortex [30], but exist only in somewhat more developed sensory areas like barrel cortex or in cat and primate V1.

Fourth, except from the well-known L5 extratelencephalic (ET) projection neurons and some characteristic morphologies in L6 (subplate and inverted cells), our data and methods suggest that excitatory neurons in the mouse visual cortex form mostly a continuum with respect to dendritic morphology. Previous studies, in contrast, work on the premise that discrete cell types exist and categorize neurons into up to 20 m-types [25, 21, 24, 10, 16, 8, 32], most of them using clustering methods on morphological features [25, 22, 24, 10]. While they assume that each cluster corresponds to a distinct m-type, they report the presence of variability within their proposed m-types. Furthermore, their visualizations of morphometrics per m-type depict further intra-class variability [10, 16, 32]. Thus, we believe that our data is consistent with previous work, but our data-driven, quantitative approach suggests that the morphological landscape of cortical excitatory neurons is better described as a continuum, with a few notable exceptions in deeper layers. This notion has also been brought up recently by transcriptomics studies, which observe continuous variation among cell types in cortex [35, 36, 11, 31, 43] as well as subcortical areas [34, 39]. Furthermore, variation within transcriptomic types found in several of the studies aligns with variation observed in other modalities [11, 31]. Scala et al. [31] suggest that neurons are organized into a small number of distinct and broad “families”, each of which exhibits substantial continuous variation among its family members. In their case, a substantial degree of morphological variation was evident among excitatory neurons of the IT type, and this variation correlated with transcriptomic variation within the type as well as the cortical depth of the neuron – resembling the gradual decrease in the width of the apical tuft with increasing cortical depth we observed. Our analysis supports the notion of broad “families” with intrinsic variation: excitatory cells can be mostly separated by layers into roughly a handful of families, each of which contains a substantial degree of variation in terms of morphology, which might also co-vary with other modalities.

This result does not rule out the possibility that there are in fact distinct types; it simply suggests that features beyond dendritic morphology need to be taken into account to clearly identify these types. For instance, the results of [32] suggest that the 5P-NP cells can be separated from other layer 5 pyramidal neurons by considering the class of interneurons that target them. It is also not guaranteed that our data-driven method identifies all relevant morphological features. Every method has (implicit or explicit) inductive biases. We tried to avoid explicit human-defined features, but by choosing a graph-based input representation we provided different inductive biases than, for instance, a voxel-based representation or one based on point clouds. However, the fact that we could reconcile known morphological features, discover novel ones and achieve good classification accuracy on an annotated subset of the data suggests that our learned embeddings indeed contain a rich and expressive representation of a neuron’s dendritic morphology.

In summary, recent studies of morphological as well as transcriptomic characteristics of cortical excitatory neurons suggest the presence of a few broad families of cell types, each exhibiting considerable intrinsic variation [11, 31, 43]. Due to this continuous variation a separation into finer cell types within these families is ambiguous. This raises the question whether it is feasible to establish a comprehensive atlas of cortical excitatory cell types. We suggest that we should rather think of the variability across cells as axes of variation, understand how these axes of variation correlate between modalities and whether they are just insignificant biological heterogeneity or indeed functionally relevant.

## 4 Methods

### 4.1 Dataset

The dataset consists of a 1.3 × 0.87 × 0.82 *mm*^3^ volume of tissue from the visual cortex of an adult P75-87 mouse, which has been densely reconstructed using serial section electron microscopy (EM) [6]. We used the subvolume 65, which covers approximately 1.3 × 0.56 × 0.82 *mm*^3^. It includes all layers of cortex and spans primary visual cortex (V1) and two higher visual areas, anterolateral area (AL) and rostrolateral area (RL). We refer to the original paper on the dataset [6] for details on the identification and morphological reconstruction of individual neurons.

### 4.2 Skeletonization and cell compartment label assignment

The EM reconstructions yielded neuronal meshes. These meshes might be incomplete or exhibit different kinds of errors including merges of other neuronal or non-neuronal compartments onto the neurons. Therefore an automatic proof-reading pipeline that resulted in neuronal skeletons was executed (companion paper; Celii et al. [4]). For the skeletal detection from the reconstructed meshes, the meshes were first downsampled to 25% of their resolution and made watertight. Then glia and nuclei meshes were identified and removed. For the remaining meshes the locations of the somata were identified using a soma detection algorithm [44]. Each neurite submesh was then skeletonized using a custom skeletonization algorithm which transformed axonal and dendritic processes into a series of line segments to obtain the skeleton (companion paper; Celii et al. [4]). For each skeleton, the highest probability axon subgraph was determined and all other non-soma nodes were labeled as dendrites. A final heuristic algorithm classifies subgraphs of dendritic nodes into compartments, such as apical trunks generally projecting from the top half of somas and with a general upward trajectory and obliques as projections off the apical trunks at an approximate 90 degree angle. For further details on the compartment label assignment, please see companion paper [4].

### 4.3 Coordinate transformations

The EM volume is not perfectly aligned. First, the pial surface is not a horizontal plane parallel to the (*x, z*)-plane, but is instead slightly tilted. Second, the thickness of the cortex varies across the volume such that the distance from pia to white matter is not constant. Without any pre-processing, an unsupervised learning algorithm would pick up these differences and, for instance, find differences of layer 6 neurons across the volume simply because in some parts of the volume they tend to be located deeper than in others and their apical dendrites that reach to layer 1 tend to be larger. Using *relative* coordinates solves such issues if pia and white matter correspond to planes (approximately) parallel to the (*x, z*)-plane. To transform our coordinate system in such standardized coordinates, we first applied a rotation about the *z*-axis of 3.5 degrees. This transformation removed the systematic rotation with respect to the native axes (Fig. A.1B). To standardize measurements across depth (*y*-axis) and to account for differential thickness of the cortex, we estimated the best linear fit for both pial surface and white matter boundary by using a set of manually placed points, which are located on a regular grid along (*x, z*) with a spacing of 25 µm. For each (*x, z*)-coordinate, the *y*-coordinate was normalized such that the pia’s *z* coordinate corresponded to the average depth of the pia and the same for the white matter. This transformation resulted in an approximation of the volume where pia and white matter boundaries are horizontal planes orthogonal to the *y* axis and parallel to the (*x, z*)-plane. Fig. A.1C shows example neurons before and after normalization. All training and subsequent analysis were performed on this pre-processed data.

### 4.4 Expert cell type labels

For a subset of the neurons in the volume experts labeled neurons according the following cell types: layer 2/3 and 4 pyramidal neurons, layer 5 near-projecting (NP), extratelencenphalic (ET) and intratelencenphalic (IT) neurons, layer 6 intratelencenphalic (IT) and cortico-thalamic (CT) neurons, Martinotti cells (MC), basket cells (BC), bipolar cells (BPC) and neurogliaform cells (NGC). Cell types were assigned based on visual inspection of individual cells taking into account morphology, synapses and connectivity, nucleus features and their (*x, y, z*)-location. All neurons were taken from one 100 µm column in the primary visual cortex (see companion paper, Schneider-Mizell et al. [32]). We did not use neurons with expert labels to train GraphDINO, but used them only for evaluation.

### 4.5 Morphological feature learning using GraphDINO

For learning morphological features in an unsupervised, purely data-driven way, we used a recently developed machine learning method called GraphDINO [42]. GraphDINO maps the skeleton graph of a neuron onto a 32-dimensional feature vector, which we colloquially refer to as the neuron’s “bar code”. For training GraphDINO, each neuron’s skeleton was represented as an undirected graph *G* = (*V, E*). *V* is the set of nodes 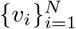 and *E* the set of undirected edges *E* ={*e*_*ij*_ = (*v*_*i*_, *v*_*j*_)} that connect two nodes *v*_*i*_, *v*_*j*_. Each node has a feature vector attached to it that holds the 3d Cartesian coordinate of the node, relative to the soma of the neuron. The soma has the coordinate (0, 0, 0), i.e. is at the origin of the coordinate system. Because axons have not been reconstructed well in the data yet, we focused on the dendritic skeleton only and removed segments labeled as axon. We trained GraphDINO on a subset of the dataset, retaining 5,113 neurons for validation and 2,941 neurons for testing. The test set was chosen to contain the 1,011 neurons that were labeled by expert anatomists into morphological cell types (Sec. 4.4; [32]), while the other 1,930 neurons were i.i.d. sampled. The training and validation sets were i.i.d. sampled from the remaining neurons with a 90%-10% split (Fig. A.4).

GraphDINO is trained by generating two “views” of the same input graph by applying random identity-preserving transformations (described below). These two views are both encoded by the same neural network. The training objective is to maximize the similarity between the embeddings of these two views. To obtain the two views of one input graph, we subsampled the graph, randomly rotated it around the *y*-axis (orthogonal to pia), dropped subbranches and perturbed node locations. When subsampling the graph, we randomly dropped all but 200 nodes, always retaining the branching points. Rotations around the y-axis were uniformly distributed around the circle. During subbranch deletion we removed *n* = 5 subbranches. For node location jittering, we used *σ* = 1. In addition, the entire graph was randomly translated with *σ* = 1. For further details on the augmentation strategies, see Weis et al. [42].

The Adjacency-conditioned Attention network architecture had seven AC-Attention layers with four attention heads each. The dimensionality of the latent representation 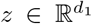 was set to *d*_1_ = 32 and the dimensionality of the projection 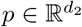 was *d*_2_ = 5, 000. All other architecture details are as described in the original paper [42]. For training, we used the Adam optimizer [18] with a batch size of 128 for 50, 000 iterations. The learning rate was linearly increased to 10^−3^ during the first 1,000 iterations and then decayed using an exponential schedule with a decay rate of 0.5.

We ran ablation experiments using different dimensionalities for the latent space *d*_1_ ∈ {16, 32, 64, 128 }and varied the number of training iterations *i* ∈ {25 000, 50 000, 100 000, 200 000 } (Fig. A.6). Additionally, we replaced the cross-entropy loss with the contrastive SimCLR loss [5] and trained variants with different mini-batch size *b* ∈ {128, 1024, 2048}(Fig. A.7), as contrastive losses have been shown to be sensitive to the number of negative samples used in the loss [5]. Training with *b* = 2, 048 diverged.

### 4.6 Morphological clustering

For qualitative inspection of the data and the analyses in Fig. 6B and Fig. 7B, we clustered the neurons using the learned vector embedding of each neuron’s morphological features. We fit a Gaussian Mixture model (GMM) with diagonal covariance matrix using scipy [27] on the whole dataset as well as per cortical layer using 60 clusters and 15 clusters, respectively. As we found no evidence that these clusters (or any other clustering with fewer or more clusters) represent distinct cell types, we did not use this clustering to define cell types, but rather think of them as modes or representing groups of neurons with similar morphological features.

### 4.7 Data quality control steps

The dataset was generated by automatic segmentation of EM images and subsequent automatic processing into skeletons. As a consequence, not all cells are reconstructed perfectly. There is a significant fraction of wrongly merged or incompletely segmented cells. We used a combination of our learned GraphDINO embeddings and supervised classifiers trained on a subset of the neurons (*n* = 1, 011) which were manually proofread and annotated by experts (see Sec. 4.4 and companion paper, Schneider-Mizell et al. [32]). Our quality control pipeline was as follows: First, we computed GraphDINO embeddings on the full dataset of 54,192 neurons (including both excitatory and inhibitory neurons). Next, we removed neurons which are close to the boundaries of the volume, as these neurons are only partly reconstructed. After this step, we were left with 43,666 neurons. Within this dataset we identified neurons which are incorrectly reconstructed using a supervised classifier described in the next section, reducing the dataset to 37,362 neurons. Subsequently, we identified interneurons using a supervised classifier described in the next section, reducing the dataset to 33,997 excitatory neurons. Finally, on this dataset we manually proofread around 480 atufted neurons. As a result, we identify and remove another set of 2,684 neurons whose reconstructions were incomplete, leaving us with a final sample size of 31,313 putative excitatory and correctly reconstructed neurons for our main analyses.

### 4.8 Supervised classifiers

To identify reconstruction errors and interneurons, we used a subset of the dataset (*n* = 1, 011) that was manually proofread and annotated with cell type labels by experts (see Sec. 4.4 and companion paper, Schneider-Mizell et al. [32]). Based on these and additional neurons we identified as segmentation errors, we trained classifiers to detect segmentation errors, inhibitory cells and cortical layer membership using our learned 32-dimensional vector embeddings of the neurons’ skeletons (see Sec. 4.5). In our subsequent analysis, we focused on neurons that were identified as complete and excitatory by our classifier. We used the inferred cortical layer labels to perform layer-specific analyses.

For all classifiers, we used ten-fold cross-validation on a grid search to find the best hyperparameters. We tested logistic regression with the following hyperparameters: type of regularization (none, L1, L2 or elastic net), regularization weight *C* ∈ 0.5, 1, 3, 5, 10, 20, 30 and whether to use class weights that are inversely proportional to class frequencies or no class weights. In addition, we tested support vector machines (SVMs) with the following hyperparameters: type of kernel (Linear, RBF or polynomial), L2 regularization weight *C* ∈ 0.5, 1, 3, 5, 10, 20, 30 and degree of polynomial *d* ∈ 2, 3, 5, 7, 10, 20 for the polynomial kernel and whether to use class weights or no weights. After having determined the optimal hyperparameters using cross-validation, we retrained the classifier using the optimal hyperparameters on its entire labeled set.

#### Removal of fragmented neurons

To remove fragmented neurons prior to analysis, we trained a classifier to differentiate between the manually proofread neurons from all layers (*n* = 1, 011) and fragmented cells (*n* = 240). We identified fragmented cells using our clustering of the vector embeddings of the whole dataset without boundary neurons (*n* = 43, 666) into 25 clusters per layer and manually identify clusters that contained fragmented cells (2–3 clusters per layer). We then sampled 60 fragmented cells per layer as training data for our classifier.

We trained a support vector machine (SVM) using cross-validation as described above. Its cross-validated accuracy was 95% (Fig. A.3A). The best hyperparameters were: polynomial kernel of degree 4 and *C* = 3. We used those hyperparameters to retrain the classifier on the full training set of 1,251 neurons. Using this classifier, we inferred whether a neuron is fragmented for the entire dataset (*n* = 43, 666). We then removed cells predicted to be fragmented (*n* = 6, 304) from subsequent analyses.

To validate the classification into fragmented and whole cells, we manually inspected ten neurons that were not in “fragmented” clusters before classification, but were flagged as fragmented by the classifier. Nine out of ten had missing segments due to segmentation errors or due to apical dendrites leaving the volume.

#### Removal of inhibitory neurons

Analogously, we trained a classifier to predict whether a neuron is excitatory or inhibitory by using the manually proofread and annotated neurons (*n* = 1, 011) (Sec. 4.4). As input features to the classifier we used our learned embeddings and additionally two morphometric features: synaptic density on apical shafts (number of synapses per micrometer of skeletal length except those located on spines) and spine density (number of spines per micrometer of skeletal length). These two features have been shown to separate excitatory from inhibitory neurons well in previous work (see companion paper, Celii et al. [4]). The annotated dataset contains 922 excitatory and 89 inhibitory neurons.

We trained a logistic regression. Its cross-validated accuracy was 99% (Fig. A.3B). The best hyperparameters were: L2 regularization (*C* = 5) and using class weights. We used those hyperparameters to retrain on the full training set of 1,011 neurons. Using this classifier, we inferred whether a neuron is excitatory or inhibitory for the entire dataset after removing fragmented cells and after the removal of 227 neurons that do not have spine and synapse densities available (*n* = 37, 135). We then removed all inhibitory cells from subsequent analyses (*n* = 3, 138).

#### Inference of cortical layers

To determine cortical layer labels for the entire dataset, we followed a two-stage procedure. First, we inferred the layer of each neuron using a trained classifier. Then we determined anatomical layer boundaries based on the optimal cortical depth that separates adjacent layers.

We first trained a SVM classifier for excitatory cells on the 922 manually annotated excitatory neurons by pooling the cell type labels per layer. Its cross-validated balanced accuracy was 89% (Fig. A.3C). The best hyperparameters were: polynomial kernel of degree 5, *C* = 3. Using this classifier, we inferred the cortical layer of all excitatory neurons (*n* = 33, 997; Fig. 2).

The spatial distribution of inferred layer assignments was overall well confined to their respective layers. As expected, there was some spatial overlap of labels at the boundaries, since layer boundaries are not sharp. We nevertheless opted for assigned neurons to layers based on their anatomical location rather than their inferred label. To do so, we determined the optimal piece-wise linear function that separated two consecutive layers. Thus, the layer assignments used for subsequent analyses were purely based on the soma depth of each neuron relative to the inferred layer boundaries – not on the classifier output.

#### Inference of coarse cell type labels

In Fig. 5 we show cell type labels for layer 5. These were determined by training a SVM to classify the excitatory neurons into cell types using the 922 manually annotated neurons. The cross-validated balanced accuracy of this classifier was 85% (Fig. A.3D). The best hyperparameters were: polynomial kernel of degree 2, *C* = 20, using class weights. Using this classifier, we inferred cell type labels for all excitatory neurons (*n* = 33, 997).

### 4.9 Manual validation of apical skeletons

We found a significant fraction of atufted neurons across layers 4–6. To determine the extent to which these cells are actually atufted or an artifact of incomplete reconstructions, we manually inspected 479 neurons in Neuroglancer [9] with respect to the validity of their apical termination. During manual inspection, we annotated neurons’ reconstruction as “naturally terminating,” “out-of-bounds,” “reconstruction issue” or “unsegmented region.” Reconstruction issues were the case where the EM slice was segmented correctly, but the tracing missed to connect two parts of the same neuron. Unsegmented regions were the case where one or multiple EM images or parts thereof were not segmented correctly and therefore the neuron could not be traced correctly. In addition, we classified the neurons as either “atufted,” “small tufted” or “tufted,” both before validation and after correcting reconstruction errors.

For layer 4, we inspected 120 atufted neurons. Of those, 64% have missing segments on their apical dendrites and 36% have a natural termination. Note, however, that 74% of the neurons had a consistent tuft before and after validation. Even though parts of the apical dendrite were missing, qualitatively the degree of tuftedness did not change. For atufted neurons this means that their apical dendrite merely terminated early, but this reconstruction error did not change their classification as atufted. In layer 4, neurons with a natural termination end more superficially than neurons with missing segments. We therefore excluded L4 neurons from the analysis whose apicals end more than 154 micrometers below the pia to exclude neurons with reconstruction errors from our analysis. This threshold was selected such that the F1-score is maximized, i.e. retaining as many atufted neurons with natural termination, while removing as many neurons with missing segments as possible. The threshold was computed on the 120 validated neurons. This process excludesd 557 neurons from layer 4.

For layer 5, we inspected 176 neurons with early-terminating apical dendrites. Of those, 59 showed a natural apical termination, while 117 had reconstruction issues or left the volume. We found no clear quantitative metric like the depth of the apical to exclude neurons with unnatural terminations. Therefore, we excluded neurons based on their cluster membership from further analysis if the cluster contained more than 50% of neurons with unnatural terminations. Of the 15 clusters, we excluded four, corresponding to 1,258 out of 5,858 L5 neurons.

For layer 6, we inspected 183 neurons with early terminating apicals. Of those, 100 showed a natural apical termination, while 83 had reconstruction issues or left the volume. Due to the slant of the volume, long, narrow L6 cells near the volume boundary have a high likelihood of leaving the boundary with their apical dendrite. Therefore, we excluded all L6 neurons whose apical dendrite left the volume (*n* = 867) prior to our analysis. We considered a neuron as leaving the volume if the most superficial point of its apical tree is within a few micrometers of the volume boundary.

Overall, we excluded 2,684 neurons as a result of this manual validation step, resulting in a final sample size of 31,313 neurons used in our analysis (Figs. 5+6+7).

### 4.10 Cortical area boundaries

Cortical area boundaries were manually drawn from retinotopic maps of visual cortex taken before EM imaging. For further details see companion paper [6].

### 4.11 Dimensionality reduction

For visualization of the learned embeddings, we reduced the dimensionality of the 32d embedding vector to 2d using t-distributed stochastic neighbor embedding (t-SNE; [37]) using the openTSNE package [28] with cosine distance and a perplexity of 30 for t-SNE plots of individual cortical layers and a perplexity of 300 for the whole dataset. The perplexity of t-SNE needs to be set dependent on the dataset size. We followed the recommendation of Kobak and Berens [19] of setting it to *perplexity* = *n/*100, which led to the approximate perplexity of 300 for our dataset of *n* = 33, 997 excitatory cells. However, to show that our interpretation is not restricted to this specific perplexity we visualized additional runs with *p* ∈ *{*30, 100, 1 000*}* (Fig. A.8). Additionally, we used UMAP [23] and PaCMAP [40] with different number of neighbors *p* ∈ *{*30, 100, 300, 1 000*}* to show that our interpretation is not dependent on the use of t-SNE (Fig. A.8).

### 4.12 Morphometric descriptors

We computed morphometrics based on the neuronal skeletons for the analysis of the learned latent space. Morphometrics were not used for learning the morphological vector embeddings. We computed morphometrics based on compartment labels: soma, apical dendrites, basal dendrites and oblique dendrites (Sec. 4.2). They are visualized in Fig. 4. Total Apical length is defined as the total length of all segments of the skeletons that are classified as apical dendrites. Total Basal length is computed analogously. Depth refers to the depth of the soma centroid relative to the pia after volume normalization (Sec. 4.3), where pia depth is equal to zero. Height is the absolute difference between the highest and the lowest skeleton node of a neuron in *y*-direction. Apical width refers to the widest extent of apical dendrites in the (*x, z*)-plane. Basal bias describes the difference between the soma depth and the center of mass of the basal dendrites along the *y*-axis. Due to the dataset size, compartment labeling was done automatically (see companion paper [4]). However, identifying apical dendrites rule-based does not work well for all neurons. For instance, it fails for the inverted L6 neurons [4]. For Fig. 5, we removed neurons for which the automatic morphometric pipeline failed. For layer 2/3 10,196 of 10,564 neurons are included in the analysis, for layer 4 7,751 of 7,775, for layer 5 4,443 of 4,600 and for layer 6 8,274 of 8,374. The GraphDINO feature space has the advantage of being independent of knowing which branches are apical and which are basal dendrites. However, our downstream analysis relies on it in certain parts (Fig. 5+Fig. 6+Fig. 7).

### 4.13 Statistics

Apical lengths in Sec. 2.8 were compared between V1 and HVA per laminar layer with four independent two-tailed Student’s t-tests. The single-test significance level of 0.01 was corrected to 0.0025 for multiple tests using Bonferroni correction. Only neurons that have any nodes labeled as apical were taken into account for this analysis. In L2/3, *n* = 6, 760 neurons were taken into account from V1 and *n* = 3, 436 from HVA; for L4 *n* = 5, 217 (V1) and *n* = 2, 534 (HVA); for L5 *n* = 3, 708 (V1) and *n* = 1, 924 (HVA); and for L6 *n* = 3, 959 (V1) and *n* = 2, 618 (HVA).

### 4.14 Cluster analysis

#### Generation of synthetic data

To obtain synthetic data distributions that are close to the neuronal data, we first fit Gaussian mixture models (GMMs) with the number of components *n* ∈ {10, 20, 40}and diagonal covariance matrices to the neuronal embeddings, extracting cluster means and weights of the fit mixture components. Using these, we subsequently generated synthetic data from Gaussian mixtures with isotropic covariance matrices with increasing variances spanning the space from distinctly separated clusters to continuous distributions (Fig. 3B & Fig. A.5).We used variances *σ*^2^ ∈ {0.005, 0.01, 0.03, 0.05, 0.07, 0.1, 0.3, 0.5, 0.7, 1.0}for each number of components *n* ∈ {10, 20, 40}, resulting in 27 synthetic datasets. For each Gaussian mixture, we drew 33,997 samples equivalent to the number of excitatory neurons. Samples were 32-dimensional like the morphological embeddings.

#### ARI analysis

To judge whether the correct number of clusters can be recovered, we split the data (both synthetic datasets and the neuronal data) into training and validation data (90%:10% split). For each synthetic dataset and the neuronal data, we fit 100 GMMs with number of components ∈ {7, 10, 15, 20, 40, 60, 80}and isotropic covariance matrix. We then computed the pairwise adjusted rank index (ARI) between the different clustering runs for the same number of components and report the average ARI on the validation set (Fig. 3B & Fig. A.5). All visualizations show clustering runs with the best log-likelihood score on the validation set (Fig. 3).

#### Unimodality versus bimodality of neighboring clusters

To examine if two neighboring clusters (neighboring in terms of least Euclidean distance between cluster means) form rather a unior bimodal distribution, we first projected the samples of the two clusters onto the line connecting the two cluster means. We then visualized the 1d histogram as well as the cumulative distribution function (CDF) of the samples from both clusters. Additionally, we computed the dip statistic [12] to quantify how close two neighboring clusters are to forming a unimodal distribution.^1^ We scaled the dip statistic with a factor of 4 such that the extreme case of two delta distributions at *x*_*i*_ and *x*_*j*_ with *i≠j* result in dip = 1. As exemplified by the synthetic data, when neighboring clusters evolve from discrete clusters to forming a continuum, the dip statistic decreases and the CDF forms a smooth curve (Fig. 3B, grey insets 1–6).

#### Connectivity graph

For each cluster of the Gaussian mixture model with 20 components of the neuronal data, we computed the dip statistic to its three nearest neighbors based on Euclidean distance in the 32-dimensional embedding space. We thresholded the neighbor selection by the average distance of all clusters to their third-nearest neighbor to avoid including spurious connections between clusters that do not have any close neighbors (threshold = 2.38 Euclidean distance in latent space). The line width of the graph (Fig. 3F) was determined as the inverse dip statistic between the nearest neighbors. Additionally, we computed the maximum dip statistic between all clusters and their nearest neighbor for the neuronal data and the synthetic datasets (Fig. 3G).

### 4.15 Prediction of morphological features from functional bar codes

The MICrONS dataset encompasses EM images as well as Calcium imaging of the same portion of the visual cortex of one mouse [6]. The companion paper by Wang et al. [38] created a digital twin of the functional properties of the neurons from the Calcium imaging data (Fig. 7C). We used the resulting functional embeddings of the neurons as input features to a linear regression model to predict the basal bias metric of the layer 4 neurons, thereby predicting a morphological feature from the functional properties of the neurons. There are 2,347 L4 neurons in V1 with both functional and morphological data available. We performed nested cross-validation to select hyperparameters and report test set performance using 10-fold cross validation for the inner and the outer loop. To select hyperparameters a grid search over regularization strength *α* ∈ {0.01, 0.1, 0.5, 1, 5, 10} as well as L1 to L2 ratio ∈ {0, 0.25, 0.5, 0.75, 1.0} was performed. The best model had a *R*^2^-score of 0.17 and ground truth and predicted basal bias had a Pearson correlation of 0.41 (Fig. 7D, *p* < 10^−10^). To control for soma depth as a confounder, we repeated the analysis predicting the basal bias from the soma depth as well as from the functional embeddings in addition to the soma depth, resulting in *R*^2^ = 0.28 for both predictors vs. 0.21 for depth only (*ρ* = 0.53, *p* < 10^−10^ and *ρ* = 0.46, *p* < 10^−10^, respectively). We tested the difference in the correlation coefficients using a two tailed Fisher’s z-test resulting in a significant difference between the two (*p* = 0.0015).

## Data availability

Data for this paper was analyzed at materialization version 374. Data is publicly available via https://www.microns-explorer.org/cortical-mm3 and will be updated closer to publication.

## Code availability

The code for GraphDINO is available at https://eckerlab.org/code/weis2021b/. Analysis code will be made available on the Eckerlab Github repository (forthcoming). Analyses were performed in Python 3.10 using custom code and the libraries Matplotlib362, Numpy124, openTSNE062, Pandas152, Pytorch113, Scikit-learn120, Scipy110, and Seaborn012 for general computation, machine learning and data visualization.

## Author Contributions

We use the CRediT system for author roles. Conceptualization: ASE, MAW, AST, PB. Methodology: MAW, PB, ASE, EYW. Software: MAW, SP, TL. Validation: MAW, SP. Formal analysis: MAW, SP, LH. Investigation: MAW, SP, BC. Resources: BC, PGF, JAB, ALB, DB, JB, DJB, MAC, FC, NMdC, SD, LE, AH, ZJ, CJ, DK, NK, SK, KiL, KaL, RL, TM, GM, EM, SSM, SM, BN, SP, RCR, CMSM, HSS, WS, MT, RT, NLT, WW, JW, WY, SY. Data curation: MAW, SP, BC. Writing - Original draft: MAW, SP, ASE. Writing - Review & editing: MAW, SP, LH, TL, AST, ASE. Visualization: MAW, SP, TL. Supervision: ASE, AST, JR. Project administration: ASE. Funding acquisition: ASE, AST, JR.

## Acknowledgements

M.A.W. was supported by the International Max Planck Research School for Intelligent Systems (IMPRS-IS), Tübingen. This work was supported by the European Research Council (ERC) under the European Union’s Horizon Europe research and innovation programme (Grant agreement No. 101041669). The work was supported by the Intelligence Advanced Research Projects Activity (IARPA) via Department of Interior/Interior Business Center (DoI/IBC) contract number D16PC00003, D16PC00004, and D16PC0005. The U.S. Government is authorized to reproduce and distribute reprints for Governmental purposes notwithstanding any copyright annotation thereon. Disclaimer: The views and conclusions contained herein are those of the authors and should not be interpreted as necessarily representing the official policies or endorsements, either expressed or implied, of IARPA, DoI/IBC, or the U.S. Government. The authors thank David Markowitz, the IARPA MICrONS Program Manager, who coordinated this work during all three phases of the MICrONS program. We thank IARPA program managers Jacob Vogelstein and David Markowitz for co-developing the MICrONS program. We thank Jennifer Wang, IARPA SETA for her assistance. We also thank the Allen Institute for Brain Science founder, Paul G. Allen, for his vision, encouragement and support. This work was also supported by the National Institute of Mental Health under Award Numbers R01 MH109556, P30EY002520, the NSF NeuroNex program through grant NSF-1707400, the National Institute of Mental Health and National Institute of Neurological Disorders And Stroke under Award Number U19MH114830, and the National Eye Institute award number R01 EY026927 as well as Core Grant for Vision Research T32-EY-002520-37.

## Footnotes

1 Dip statistic was computed using the python package *diptest* (https://pypi.org/project/diptest/).

