## Appendix for "An unsupervised map of excitatory neurons’ dendritic morphology in the mouse visual cortex"

[1]Institute of Computer Science and Campus Institute Data Science, University of Göttingen, Germany
[2]Institute for Theoretical Physics, University of Tübingen, Germany
[3]Max Planck Institute for Dynamics and Self-Organization, Göttingen, Germany
[4]Department of Neuroscience, Baylor College of Medicine, Houston, USA
[5]Center for Neuroscience and AI, Baylor College of Medicine, Houston, USA
[6]Princeton Neuroscience Institute, Princeton University, USA
[7]Department of Computer Science, Princeton University, USA
[8]Department of Electrical Engineering, Princeton University, USA
[9]Allen Institute for Brain Science, Seattle, WA, USA
[10]Rice University, Houston, TX, USA
[11]Massachusetts Institute of Technology, Cambridge, MA, USA
[12]Hertie Institute for AI in Brain Health, University of Tübingen, Germany
[13]Tübingen AI Center, Tübingen, Germany
[14]Department of Ophthalmology, Stanford University, Stanford, CA, USA
[15]Department of Electrical Engineering, Stanford University, Stanford, CA, USA
[16]Stanford BioX, Stanford University, Stanford, CA, USA
[17]Wu Tsai Neurosciences Institute, Stanford University, Stanford, CA, USA

#### Abstract

Neurons in the neocortex exhibit astonishing morphological diversity which is critical for properly wiring neural circuits and giving neurons their functional properties. However, the organizational principles underlying this morphological diversity remain an open question. Here, we took a data-driven approach using graph-based machine learning methods to obtain a low-dimensional morphological "bar code" describing more than 30,000 excitatory neurons in mouse visual areas V1, AL and RL that were reconstructed from the millimeter scale MICrONS serial-section electron microscopy volume. Contrary to previous classifications into discrete morphological types (m-types), our data-driven approach suggests that the morphological landscape of cortical excitatory neurons is better described as a continuum, with a few notable exceptions in layers 5 and 6. Dendritic morphologies in layers 2–3 exhibited a trend towards a decreasing width of the dendritic arbor and a smaller tuft with increasing cortical depth. Inter-area differences were most evident in layer 4, where V1 contained more atufted neurons than higher visual areas. Moreover, we discovered neurons in V1 on the border to layer 5 which avoided deeper layers with their dendrites. In summary, we suggest that excitatory neurons' morphological diversity is better understood by considering axes of variation than using distinct m-types.

### A Appendix

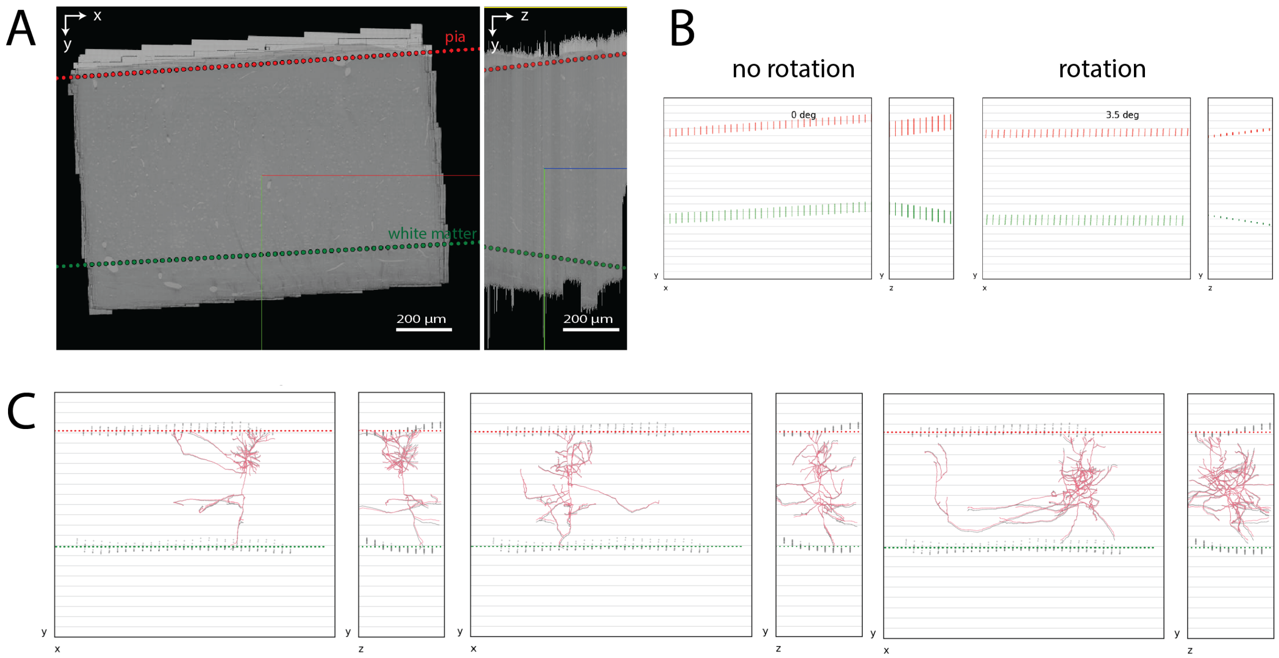


Figure A.1: **Volume pre-processing. A.** $xy$- (left) and $yz$- (right) 2D cross-sectional views of the EM volume as seen in Neuroglancer. Red scatter points - linear model of pia, Green scatter points - linear model of L6 - white matter boundary. **B.** Pia and white matter boundary models shown with (right) and without (left) rotating the volume by 3.5 degrees about the $z$-axis. **C.** Three example excitatory neuronal skeletons shown from two 2D projections ($xy$) and ($yz$) after rotation and depth normalization to the mean pia and white matter depths. Red scatter points - pia model after normalization. Green scatter points - white matter boundary after normalization. Gray shadow - pia, white matter and neuronal skeleton after rotation but before normalization.

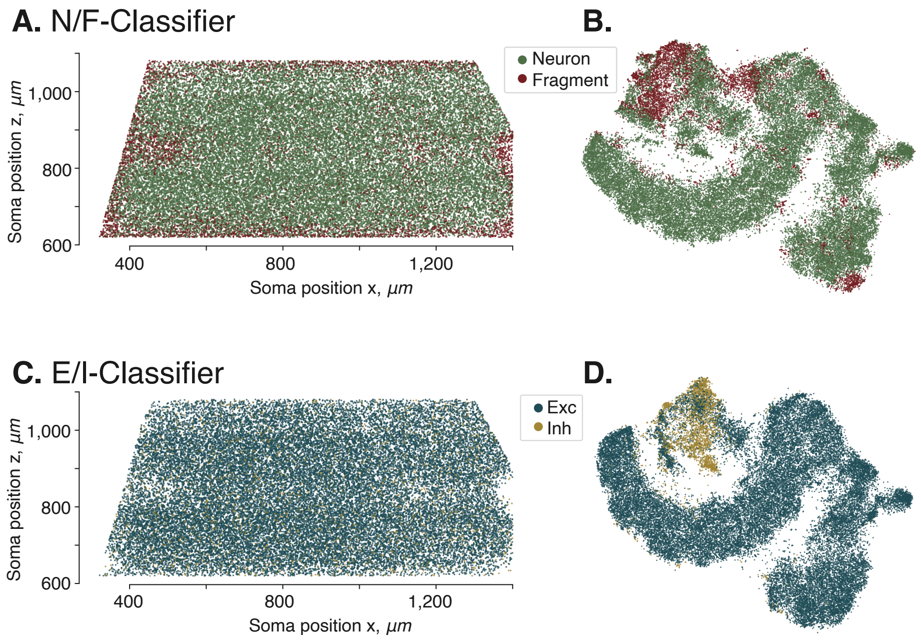


Figure A.2: **A.** Top view of the volume showing the distribution of complete neuron versus fragments predictions by our classifier based on the learned morphological embeddings. Density of fragmented neurons is high at the volume borders since neurons have a high likelihood of leaving the imaged volume with their dendrites. Additionally, we see a high number of fragmented neurons in areas where we know there have been issues during the imaging process, proving that the classifier works as intended. **B.** t-SNE embedding of neuronal morphologies colored by neuron-fragment predictions. **C.** Top view of the volume showing a uniform distribution of excitatory versus inhibitory neurons across the volume. **D.** t-SNE embedding of neuronal morphologies colored by excitatory-inhibitory predictions.

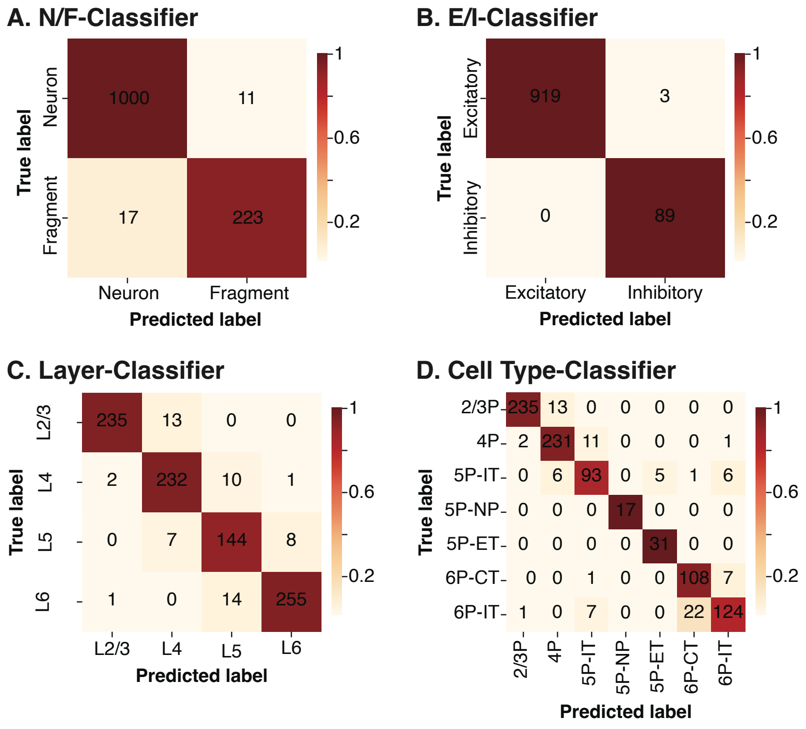


Figure A.3: Confusion matrices of classifiers trained on the full set of neurons with labels. Colors represent normalized confusion values across rows, while numerical values denote absolute confusion counts.

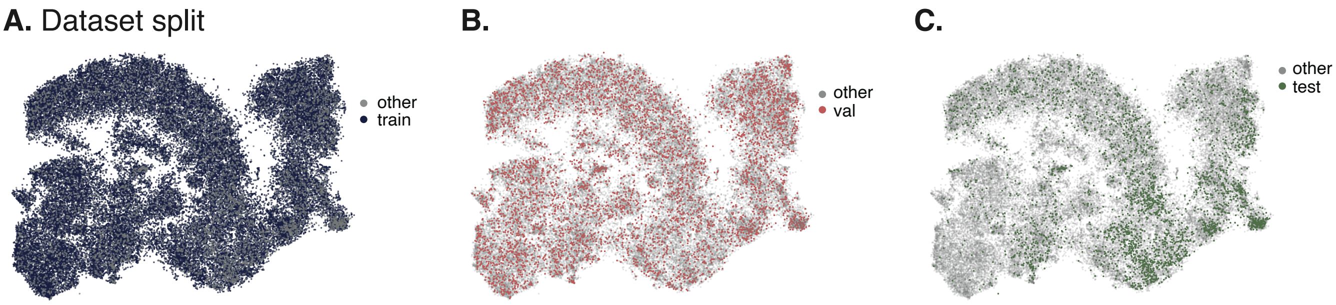


Figure A.4: t-SNE embedding of neuronal morphologies colored by the dataset split: **A.** training, **B.** validation and **C.** test set as used for GraphDINO training.

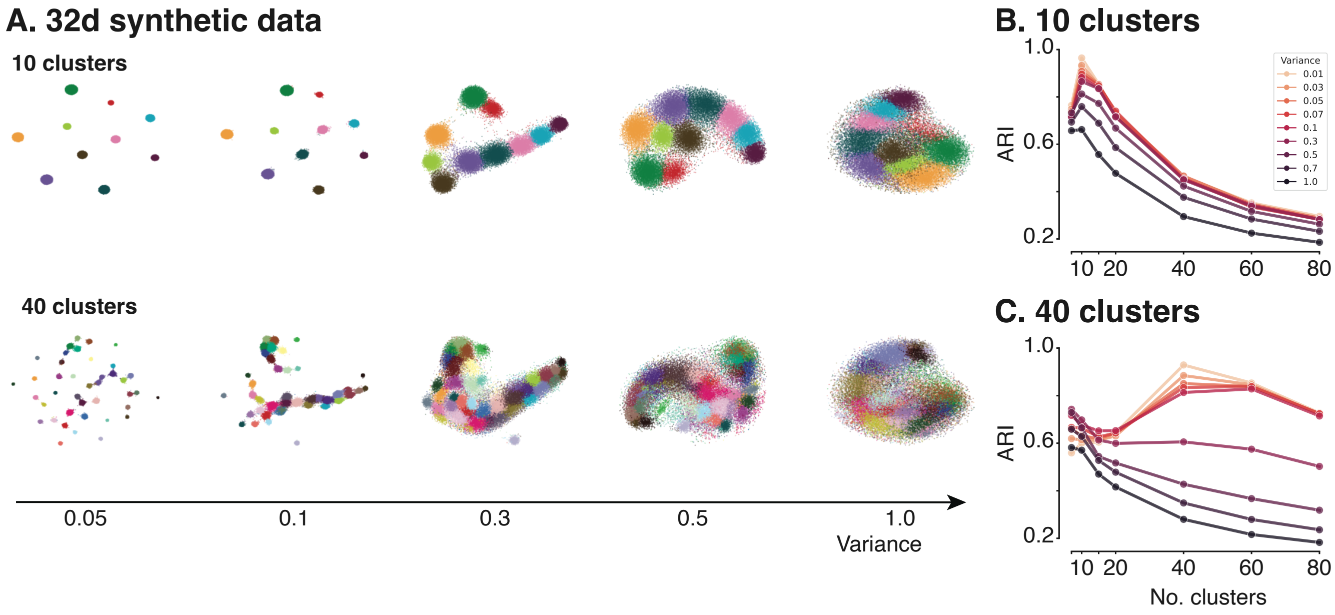


Figure A.5: **A.** t-SNE representation of synthetic data ($n = 33\,997$, perplexity= 300). Synthetic data is sampled from Gaussian mixtures with 10 (first row) and 40 components (second row). Cluster means and weights are estimated from neuronal data. Isotropic variance is set to obtain data evolving from discrete clusters to uniform distributions (left to right). **B.** Mean adjusted rand index (ARI) of 100 GMMs with increasing number of components fit to the synthetic datasets with 10 components. **C.** as **B.** but for synthetic datasets with 40 components. The correct number of underlying components can be identified as long as the variance in the data is not too high ($\sigma^2 < 1.0$ for 10 components and $\sigma^2 < 0.07$ for 40 components).

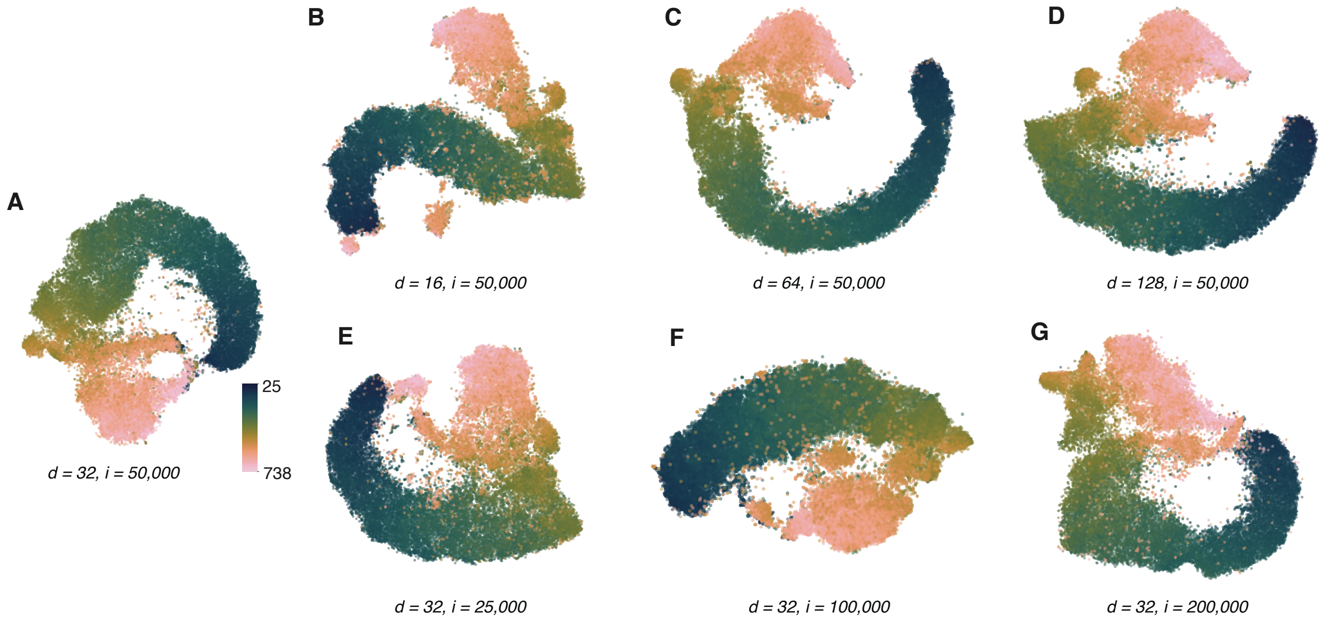


Figure A.6: t-SNE representation (perplexity=300) of morphological embeddings obtained by training GRAPHDINO with different hyper-parameters ($d$: latent dimensionality of the morphological embeddings; $i$: training iterations). Colors reflect soma depth.

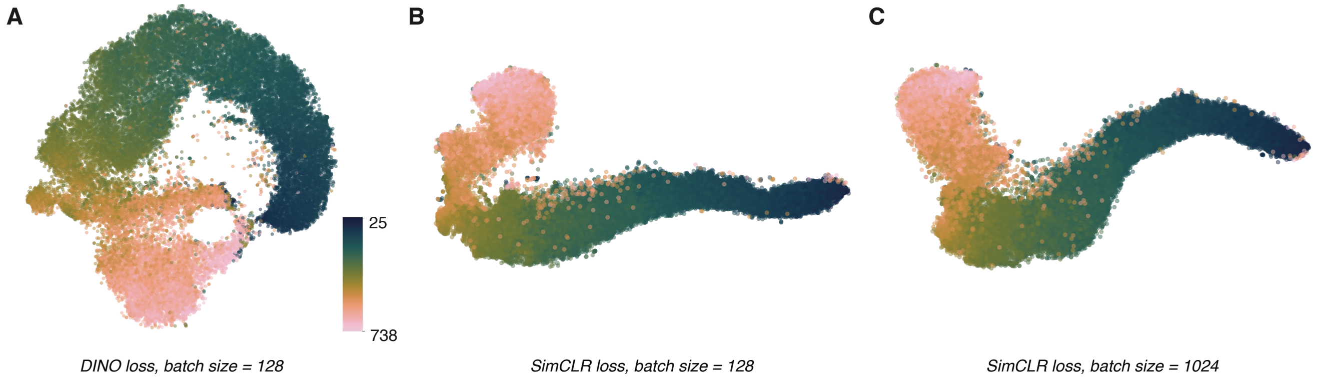


Figure A.7: t-SNE representation (perplexity=300) of morphological embeddings obtained by training GRAPHDINO with DINO loss [3] compared to SIMCLR loss [5]. Colors reflect soma depth.

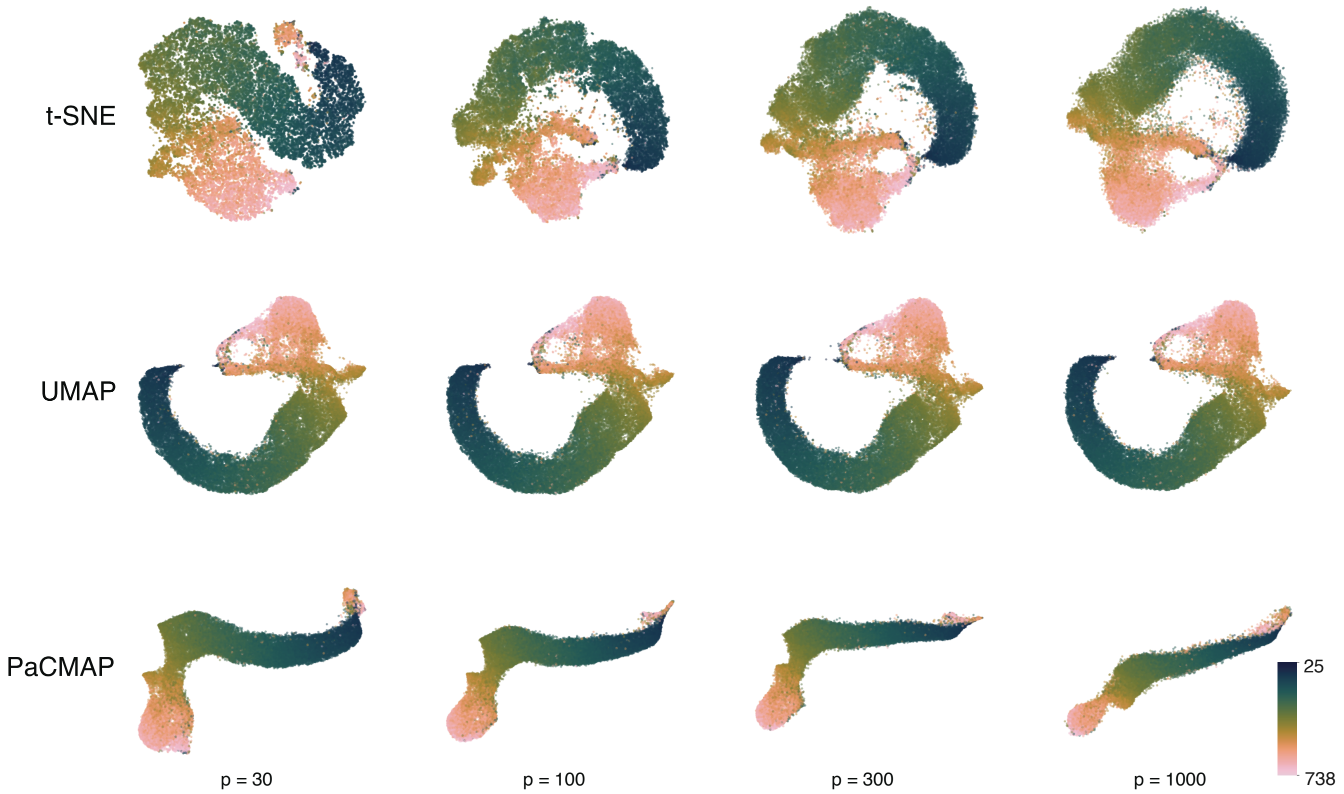


Figure A.8: **Different dimensionality reduction techniques. A.** t-SNE [37], $p$ corresponds to perplexity. **B.** UMAP [23], $p$ corresponds to number of neighbors. **C.** PaCMAP [40], $p$ corresponds to number of neighbors.

| **Apical dendrites (AD)** | **Basal dendrites (BD)** | **Others** |
|---|---|---|
| Bias x | Bias x | Soma depth |
| Bias z | Bias z | AD fraction above BD |
| Extent x | Extent x | AD fraction below BD |
| Extent z | Extent z | AD fraction of intersection with BD |
| Mean contraction | Mean contraction | AD EMD with BD |
| Maximum branch order | Maximum branch order | BD fraction above AD |
| Early branch path | Maximum Euclidean distance | BD fraction below AD |
| Number of branches | Number of branches | BD fraction of intersection with AD |
| Number of outer bifurcations | Number of stems | |
| Soma percentile x | Soma percentile x | |
| Soma percentile z | Soma percentile z | |
| Total length | Total length | |
| Depth PC 0 | | |
| Depth PC 1 | | |
| Depth PC 2 | | |
| Depth PC 3 | | |

Table A.1: List of 36 morphometrics used for analyses in Sec. A.1 (Fig. A.9A,D).

#### A.1 Comparison to previous morphological feature spaces

To compare our learned embedding space to previous morphological feature spaces, we extracted morphometrics for two subsets of the MICrONS data: for all excitatory neurons and for the "column" subset. First, we computed 36 morphometrics akin to those proposed by [10] for cell type identification of cortical neurons for all excitatory neurons after deploying the above described quality control steps. This resulted in a morphometric description of 28,312 excitatory neurons that possess the necessary annotation of apical and basal dendrites [4]. The complete list of morphometrics used can be found in table A.1. The normalized morphometrics were computed using the SKELETON-KEYS package[2] after volume normalization (Sec. 4.3). For a description of the individual morphometrics, see Gouwens et al. [10].

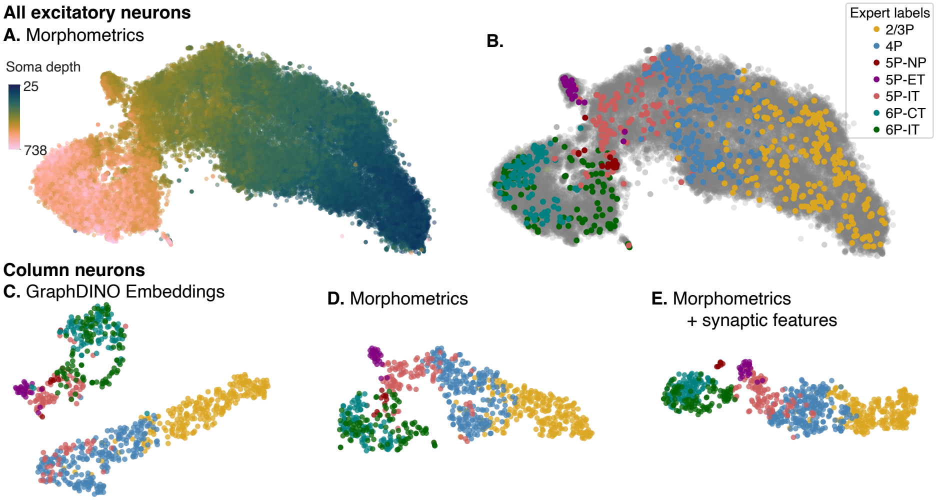


Figure A.9: t-SNE representation ($n = 28\,312$, perplexity $= 300$) of morphometric features akin to Gouwens et al. [10] for the excitatory neurons of the MICrONS dataset colored by soma depth (**A.**) and colored by expert cell types if available otherwise colored in grey (**B.**). t-SNE representation ($n = 794$, perplexity$= 30$) of features for the "column" dataset using GRAPHDINO embeddings (**C.**), morphometric features (**D.**) and morphometric + synaptic features (**E.**).

Qualitatively, we found that using explicit morphometric features instead of our learned feature embeddings leads to a feature embedding of the neuronal skeletons that is similarly or even more continuously varying along the depth of the neurons (Fig. A.9A and Fig. A.9D). Equivalent to the GRAPHDINO embeddings, the L5-ET neurons are separated from the continuum (Fig. A.9B).

One caveat here is that the morphometrics rely on the correct labeling of apical and basal dendrites, which has been done manually in the Gouwens et al. [10] study. This is not feasible for the dataset size of MICrONS. We therefore rely on automatic compartment labeling (see companion paper [4]), which is known to not work well for L2 and L6 cells (inverted, horizontal) [4]. In contrast, the GRAPHDINO feature space is independent of knowing which branches are apical and which are basal dendrites.

Second, we performed a quantitative comparison between our learned features, the morphometric features [10] and the morphometric features with additional synaptic features as used by the companion paper by Schneider-Mizell et al. [32]. The features required for the latter are only available for the hand-curated subset of the dataset (the "column" dataset) of approximately 1,000 neurons. We therefore compared the three feature spaces on this subset of the data which additionally gives us access to expert-defined cell types and therefore allows for a quantitative comparison.

When only looking at the "column" dataset, qualitatively, again all three embeddings spaces show a continuous variation (Fig. A.9C–E), especially in layers 2–4. The synaptic features are beneficial to separate L5-NP cells (Fig. A.9E) (see also [32]), while the morphometric features seem mostly to cluster the L5-ET cells (Fig. A.9D). IT neurons of L2/3 and L4 are continuously represented in all three feature spaces.

To asses our learned feature embedding quantitatively, we computed the balanced accuracy of predicting the expert cell types labels on a subset of the cells for which labels are available ($n = 794$). To this end, we trained a logistic regression model using nested cross-validation to fit hyperparameters and report test set accuracy of three different features sets: *(1)* our learned GRAPHDINO embeddings (32 dimensional), *(2)* morphometric features akin to the ones proposed by Gouwens et al. [10] (36 dimensional) and *(3)* morphometric in addition to synaptic features as used by Schneider-Mizell et al. [32] (30 dimensional: 15 morphometric and 15 synaptic features) (Tab. A.2). Note that Schneider-Mizell et al. [32] used a different set of morphometrics compared to Gouwens et al. [10]. See Tab. A.1 and the companion paper [32] for the full list of the

[2] https://skeleton-keys.readthedocs.io/

Table A.2: Balanced accuracy for expert cell type prediction from different input features using logistic regression.

| **Input Features** | **Balanced Accuracy [%]** |
|---|---|
| Morphometric features including soma depth [10] | 86.3 |
| Morphometric + synaptic features including soma depth [32] | 90.5 |
| GRAPHDINO embeddings | 84.5 |
| GRAPHDINO embeddings + soma depth | 86.4 |
| GRAPHDINO embeddings + soma depth + synaptic features | 90.5 |

morphometric features included in the feature sets. We normalized the features from Schneider-Mizell et al. [32] by subtracting the mean and dividing by the standard deviation. Without normalization both the t-SNE and the classifier accuracy were worse (56.9% balanced accuracy compared to 90.5% after normalization).

For cell type prediction, we ran nested cross-validation of a logistic regression model using ten folds for both the inner and the outer loop of the cross-validation. In the inner loop, the best hyperparameters were picked based on the validation performance. In the outer loop, the test set accuracy was determined. We report the average balanced accuracy over three trials. Hyperparameters were selected using a grid search over penalty $\in$ {None, L1, L2, ElasticNet}, regularization strength $C \in \{0.5, 1, 3, 5, 10, 20, 30\}$ and either using class weights or not.

Quantitatively, using both morphometric as well as synaptic features performed best with a balanced accuracy of 90.5%, while only using morphometric or learned features achieved 86.3% and 84.5%, respectively (Tab. A.2). Note that this is by no means a fair comparison. The cell types are defined on features, which are also reflected in the morphometric and synaptic feature sets. These feature sets were designed to discriminate between cell types manually defined by experts while the learned feature space is unsupervised. Additionally, the morphometric and synaptic feature spaces include information on anatomical depth of the neurons as well as on labeled neuronal compartments of apical and basal dendrites – information not accessible to GRAPHDINO.

When adding information on cortical depth and synapses to our learned embeddings, discriminative power was on par with morphometric features (86.4 % GRAPHDINO + depth; 90.5% GRAPHDINO + depth + synaptic features; Tab. A.2).

It is important to keep in mind, however, that the expert labels should not be considered ground truth, but are only an approximation of the current state of knowledge. It might not be possible to match individual neurons to a clear type. Furthermore, some types are rather defined by soma depth (i.e. layer 4 pyramidal cells) than by morphological properties. Therefore, interpretations of the classes need to be treated with caution.

Overall, these results show that our learned embeddings perform competitively on the task of expert label prediction with less information and despite not being specifically optimized for it. Furthermore, the interpretation of a continuum is not unique to our embeddings but is the case for other feature embeddings as well.

| Morphometric | Layer 2/3 | Layer 4 | Layer 5 | Layer 6 |
|---|---|---|---|---|
| **Depth** | 0.93 | 0.80 | 0.81 | 0.65 |
| **Height** | 0.94 | 0.81 | 0.89 | 0.93 |
| **Apical length** | 0.58 | 0.60 | 0.69 | 0.65 |
| **Apical width** | 0.59 | 0.32 | 0.44 | 0.35 |
| **Basal length** | 0.70 | 0.74 | 0.83 | 0.75 |
| **Basal bias** | 0.68 | 0.79 | 0.79 | 0.78 |

Table A.3: $R^2$ scores for regression of the morphometrics from the 32-dimensional latent embedding.

| | Depth | Height | Apical length | Apical width | Basal length | Basal bias |
|---|---|---|---|---|---|---|
| **Layer 2/3** | | | | | | |
| **Depth** | 1.0 | | | | | |
| **Height** | 0.93 | 1.0 | | | | |
| **Apical length** | -0.23 | -0.22 | 1.0 | | | |
| **Apical width** | -0.55 | -0.5 | 0.61 | 1.0 | | |
| **Basal length** | -0.15 | -0.1 | 0.19 | 0.07 | 1.0 | |
| **Basal bias** | 0.38 | 0.49 | 0.00 | -0.12 | -0.18 | 1.0 |
| **Layer 4** | | | | | | |
| **Depth** | 1.0 | | | | | |
| **Height** | 0.49 | 1.0 | | | | |
| **Apical length** | 0.00 | 0.14 | 1.0 | | | |
| **Apical width** | 0.13 | 0.1 | 0.64 | 1.0 | | |
| **Basal length** | -0.11 | -0.03 | -0.04 | -0.01 | 1.0 | |
| **Basal bias** | -0.29 | 0.18 | -0.09 | -0.11 | -0.01 | 1.0 |
| **Layer 5** | | | | | | |
| **Depth** | 1.0 | | | | | |
| **Height** | 0.81 | 1.0 | | | | |
| **Apical length** | 0.20 | 0.28 | 1.0 | | | |
| **Apical width** | 0.11 | 0.20 | 0.68 | 1.0 | | |
| **Basal length** | 0.06 | 0.30 | 0.43 | 0.37 | 1.0 | |
| **Basal bias** | 0.27 | 0.52 | 0.01 | 0.06 | 0.23 | 1.0 |
| **Layer 6** | | | | | | |
| **Depth** | 1.0 | | | | | |
| **Height** | -0.13 | 1.0 | | | | |
| **Apical length** | -0.15 | 0.53 | 1.0 | | | |
| **Apical width** | -0.20 | 0.27 | 0.72 | 1.0 | | |
| **Basal length** | -0.22 | -0.21 | -0.44 | -0.29 | 1.0 | |
| **Basal bias** | 0.66 | -0.02 | -0.17 | -0.33 | -0.08 | 1.0 |

Table A.4: Layer-wise Spearman’s rank correlation coefficient between morphometrics.

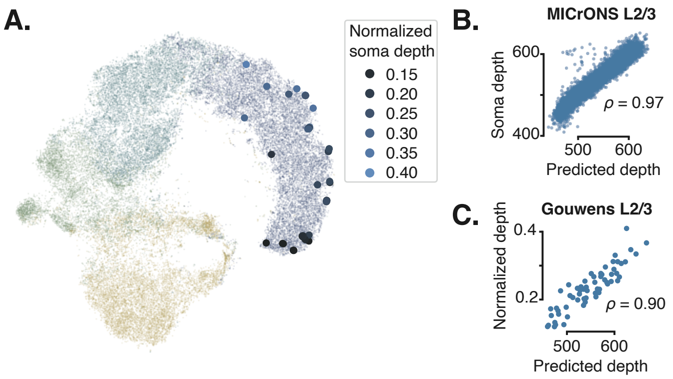


Figure A.10: **A.** t-SNE embedding of MICRONS dataset in the background ($n = 33\,997$; perplexity= 300) with Berg et al. [1] L2/3 neurons embedded in blue ($n = 61$) colored by normalized soma depth. **B.** Soma depth prediction from MICRONS latent embeddings with Pearson's correlation coefficient of $\rho = 0.97$ to anatomical soma depth. **C.** Soma depth prediction from Berg latent embeddings with Pearson's correlation coefficient of $\rho = 0.90$ to normalized anatomical depth.

#### A.2 Transfer to PatchSeq dataset

To prove the utility of our learned embedding space beyond EM datasets, we transferred our model trained on the MICRONS data to a dataset that was recorded using a different recording technique, namely Patch-Seq [2]. The Berg dataset [1] includes 136 neurons from mouse visual cortex layer 2/3 of which 61 have publicly available reconstructed dendritic morphologies. To embed the neurons in our learned feature space, we normalized the morphologies spatially along the pia–white matter axis to match the distance between pia and white matter in the MICRONS volume. To this end, we computed the first and 99th percentile of the minimum and maximum $y$-extent of the layer 2/3 neurons of the MICRONS dataset and the Berg dataset. We then computed a single scaling factor $t$ for the $y$-axis of the Berg data to match the range of the MICRONS dataset ($t = 0.927$). Subsequently, we normalized all neurons such that the soma node is centered on the origin $(0, 0, 0)$ and computed the morphological embedding for each cell by encoding it using the pre-trained GRAPHDINO model. Despite the difference in the underlying data acquisition, GRAPHDINO generalizes to this data (Fig. A.10).

To show that the feature embedding is meaningful, we trained a linear regression model to predict the soma depth from the learned embeddings of the MICRONS L2/3 neurons (Fig. A.10B; $n = 10\,564$; Pearson's correlation coefficient $\rho = 0.97$) and applied the trained linear regression model to the latent embeddings of the Berg neurons. We then calculated the Pearson correlation between the predicted depth and the ground truth normalized depth of the Berg neurons (Fig. A.10C; Pearson's correlation coefficient $\rho = 0.90$), showing that the embeddings inferred for the Berg data encode meaningful information about the underlying neurons.
